# YTHDC1 cooperates with the THO complex to prevent RNA damage-induced DNA breaks

**DOI:** 10.1101/2024.03.14.585107

**Authors:** Ning Tsao, Jennifer Olabode, Rebecca Rodell, Hua Sun, Joshua R. Brickner, Miaw-Sheue Tsai, Elizabeth A. Pollina, Chun-Kan Chen, Nima Mosammaparast

## Abstract

Certain environmental toxins are nucleic acid damaging agents, as are many chemotherapeutics used for cancer therapy. These agents induce various adducts in DNA as well as RNA. Indeed, most of the nucleic acid adducts (>90%) formed due to these chemicals, such as alkylating agents, occur in RNA^1^. However, compared to the well-studied mechanisms for DNA alkylation repair, the biological consequences of RNA damage are largely unexplored. Here, we demonstrate that RNA damage can directly result in loss of genome integrity. Specifically, we show that a human YTH domain-containing protein, YTHDC1, regulates alkylation damage responses in association with the THO complex (THOC)^2^. In addition to its established binding to *N*6-methyladenosine (m6A)-containing RNAs, YTHDC1 binds to *N*1-methyladenosine (m1A)-containing RNAs upon alkylation. In the absence of YTHDC1, alkylation damage results in increased alkylation damage sensitivity and DNA breaks. Such phenotypes are fully attributable to RNA damage, since an RNA-specific dealkylase can rescue these phenotypes. These <u>R</u>NA <u>d</u>amage-induced DNA <u>b</u>reaks (RDIBs) depend on R-loop formation, which in turn are processed by factors involved in transcription-coupled nucleotide excision repair. Strikingly, in the absence of YTHDC1 or THOC, an RNA m1A methyltransferase targeted to the nucleus is sufficient to induce DNA breaks. Our results uncover a unique role for YTHDC1-THOC in base damage responses by preventing RDIBs, providing definitive evidence for how damaged RNAs can impact genomic integrity.

## Main section

An increasing body of work in the last two decades has shown that RNAs play an important role in various DNA repair processes^3, 4^. Previous studies have revealed a role for the activating signal cointegrator complex (ASCC) in base damage repair, particularly in response to alkylation and UV damage^5–8^. More recently, we have demonstrated that RNA alkylation, and not DNA alkylation, serves as an initial signal to trigger to recruit the ALKBH3-ASCC complex^9^. The fact that a DNA repair pathway can be activated by damaged RNAs suggests that cells have evolved proper machinery to sense such RNA adducts, which could contribute to cellular damage responses. Besides being induced by chemical alkylation, RNA methylation can be generated physiologically in mammalian cells, with the most abundant RNA methylation mark being m6A^10, 11^. The YTH domain-containing proteins have been shown to recognize m6A-containing RNAs via their YTH domains to regulate various RNA functions^12^. In addition, previous studies have reported that YTH domains can also bind *in vitro* to m1A, a modification created by chemical alkylators^13, 14^. However, whether this binding is relevant or has functional consequences *in vivo* is unknown.

### YTHDC1 and the THO complex regulate alkylation responses

We reasoned that one or more YTH proteins may be involved in alkylation responses via recognition of methylated RNA. To test this, we performed an shRNA screen individually targeting all known human YTH proteins (Extended Data Fig. 1a). Previous studies have shown that the ASCC complex forms damage foci in response to alkylation^6^; thus, we monitored HA-ASCC2 foci formation as a readout of this damage response. After treating with the alkylating agent methyl methanesulfonate (MMS), we found that loss of YTHDC1 with two independent shRNAs significantly increased ASCC2 foci formation, but little effect was seen with loss of other YTH domain-containing proteins (Fig. 1a-b and Extended Data Fig. 1b). Similarly, YTHDC1 depletion also showed an elevation of endogenous ASCC3 foci upon alkylation stress (Extended Data Fig. 1c). ASCC foci form primarily in the G1 phase of cell cycle^6^; we found that depleting YTHDC1 did not affect cell cycle distribution (Extended Data Fig. 1d), indicating that the effect of YTHDC1 on ASCC foci is not due to cell cycle perturbation. Together, these results suggested that YTHDC1 may regulate a parallel alkylation response pathway.

**Fig. 1.**
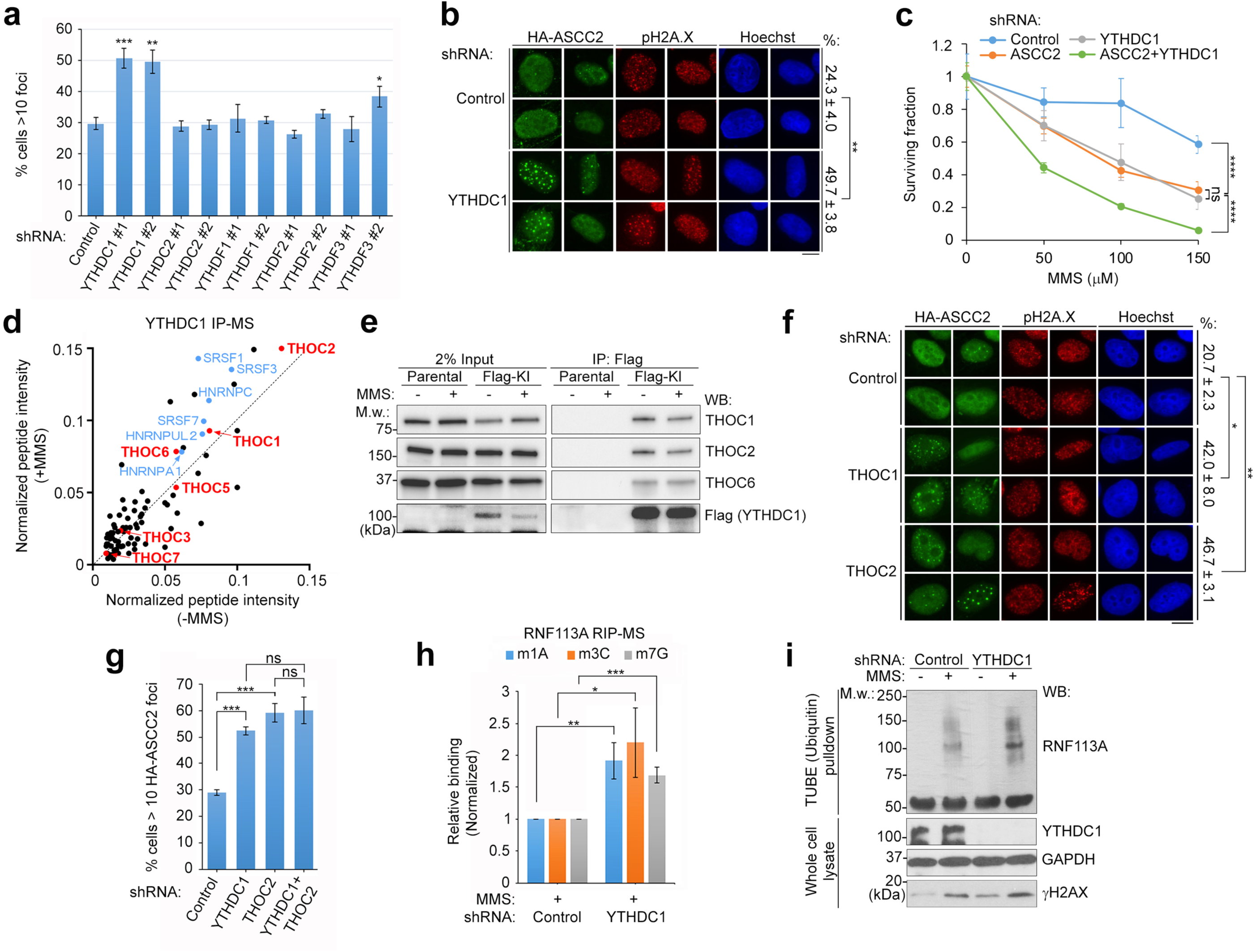
YTHDC1 and THO complex intersect with the ASCC pathway during alkylation damage. **a**, U2OS cells expressing HA-ASCC2 were infected with the indicated lentiviral shRNAs for 48 h. Cells were treated with MMS and processed for immunofluorescence staining with HA antibody, and quantified for HA-ASCC2 foci formation (n = 3 technical replicates per shRNA, at least 100 cells were analyzed in each sample; error bars indicate ± SD of the mean; *p < 0.05, **p < 0.01, ***p < 0.001 by Student’s t test). **b**, U2OS cells expressing HA-ASCC2 were infected with lentiviral control or YTHDC1 (#1 shown in **a**) shRNAs and treated with MMS, then processed for immunofluorescence staining with indicated antibodies. Scale bar, 10 µm. % cells with more than 10 HA-ASCC2 foci are shown on the right of the representative images as the mean ± SD (n = 3 biological repeats, at least 100 cells were analyzed in each sample; **p < 0.01 by Student’s t test). **c**, HeLa cells expressing inducible YTHDC1 shRNA were infected with control and ASCC2 lentiviral shRNAs with or without the induction of YTHDC1 shRNA by 0.5 µg/mL of doxycycline, then processed with MTS assay after treating with indicated concentration of MMS (n = 5 technical repeats for each sample). ****p < 0.0001 by two-way ANOVA test; ns, non-significant. **d**, Scatterplot of the endogenously Flag-tagged YTHDC1-interacting proteins identified from HeLa-S cells with or without MMS. Total peptide intensities for each were normalized with YTHDC1. Blue dots indicate the representative YTHDC1-interacting proteins identified previously^15–19^. Red dots indicate THO complex components. **e**, Parental or endogenously Flag-tagged YTHDC1 (Flag-KI) HeLa-S cell lysates were used for co-immunoprecipitation with anti-Flag (M2) resin. Immunoprecipitated (IP) and input material were analyzed by Western blotting with the indicated antibodies. **f**, U2OS cells expressing HA-ASCC2 were infected with lentiviral control, THOC1 or THOC2 shRNAs and treated with MMS, then processed for immunofluorescence staining as in **b**. Scale bar, 10 µm. % cells with more than 10 HA-ASCC2 foci are shown on the right of the representative images as the mean ± SD (n = 3 biological repeats, at least 100 cells were analyzed in each sample; *p < 0.05, **p < 0.01 by Student’s t test). **g**, U2OS cells expressing HA-ASCC2 were infected with indicated lentiviral shRNAs. Cells were treated with MMS and then processed as in **b**. Cells with more than 10 HA-ASCC2 foci were quantitated (n = 3 biological repeats, at least 100 cells were analyzed in each sample; error bars indicate ± SD of the mean; ***p < 0.001 by Student’s t test; ns, non-significant). **h**, Flag-RNF113A expressing HeLa-S cells were infected with control or YTHDC1 lentiviral shRNAs. Cells were treated with MMS followed by RNA immunoprecipitation (RIP) with anti-Flag (M2) resin. Purified RNAs were digested, dephosphorylated, and the methylated nucleosides were analyzed by LC-MS/MS. The result is shown as the mean ± SD (n = 3 independent experiments; *p < 0.05, **p < 0.01, ***p < 0.001 by Student’s t test). **i**, Control and YTHDC1 knockdown (inducible) HeLa-S cells with or without MMS treatment were harvested for the tandem ubiquitin binding element (TUBE) pulldown assay following by Western blotting with indicated antibodies (representative of three independent experiments).

Since ASCC is important for cell survival after alkylation stress, we tested the epistatic relationship of YTHDC1 and ASCC loss in alkylation damage sensitivity. While depleting either YTHDC1 or ASCC2 sensitized cells to MMS, depleting both YTHDC1 and ASCC2 showed an addictive effect on cell viability (Fig. 1c and Extended Data Fig. 1e-h). To determine the potential mechanism of YTHDC1 in this response, we identified YTHDC1 interacting proteins in HeLa-S cells expressing an endogenously Flag-tagged YTHDC1 (Extended Data Fig. 2a-c and Extended Data Table 1). Gene Ontology analysis showed that most YTHDC1 interacting partners function in RNA metabolism, consistent with previous reports^15–19^ (Extended Data Fig. 2d). We found that YTHDC1 interacted with all the THO complex members both in untreated and MMS-treated cells (Fig. 1d-e). Since the THO complex and YTHDC1 are known to promote pre-mRNA processing and export, we reasoned that they may function together in the damage response. Similar to YTHDC1 depletion, depletion of either of two THO complex members, THOC1 and THOC2, also increased ASCC2 foci formation, yet the effects were not additive when depleting both YTHDC1 and THOC2 (Fig. 1f-g and Extended Data Fig. 2e-f). Thus, YTHDC1 and the THO complex function epistatically in the alkylation response.

The fact that YTHDC1 suppressed MMS-induced ASCC foci formation suggested that YTHDC1 may compete with ASCC for the recognition of RNA methylation, which serves as the initial signal for ASCC recruitment^9^. We focused on the E3 ligase RNF113A, which is an upstream regulator of the ASCC pathway^6^. We previously showed that RNF113A associates with alkylated RNAs, resulting in RNF113A activation and autoubiquitination^6, 9^. First, we tested the binding of RNF113A to methylated RNAs upon YTHDC1 depletion using RNA immunoprecipitation coupled with quantitative mass spectrometry (RIP-MS)^9, 20^. This showed that the interaction between RNF113A and methylated RNAs is significantly increased when YTHDC1 is lost (Fig. 1h and Extended Data Fig. 2g). Furthermore, we found that autoubiquitination of endogenous RNF113A induced by MMS was further enhanced in cells that lacked YTHDC1 (Fig. 1i). Taken together, these results indicated that YTHDC1 antagonizes the activation of ASCC induced by chemical alkylation, potentially by competing with methylated RNAs.

### YTHDC1 binds to m1A-containing RNAs upon alkylation

In addition to binding m6A, previous *in vitro* studies have suggested that YTH proteins may also bind m1A, a major RNA base adduct induced by methylating agents^21, 22^. To test YTHDC1 binding to m1A-containing RNAs in cells, we performed RIP-MS with Flag-YTHDC1 expressing cells. The result showed that YTHDC1 primarily bound to m6A under control conditions (Fig. 2a and Extended Data Fig. 3a). After MMS treatment, the interaction between YTHDC1 and m6A was unchanged yet binding to m1A was significantly increased (Fig. 2a). It was possible that the association between YTHDC1 and m1A shown by RIP-MS could be a consequence of binding to another methylation mark in the same RNA which contains m1A modification. To further demonstrate the specificity of m1A binding by YTHDC1, we targeted the m1A RNA methyltransferase TRMT61A to a single locus reporter system^9, 23, 24^ (Fig. 2b). The expression of the LacI-fused methyltransferase generated the m1A modification at the site where it is targeted (Extended Data Fig. 3b). By co-staining with endogenous YTHDC1, we found that the wildtype, but not catalytically inactive (D181A) TRMT61A^9, 25, 26^ recruited YTHDC1 (Fig. 2b). This result further demonstrated that YTHDC1 can be recruited to sites containing m1A methylation.

**Fig. 2.**
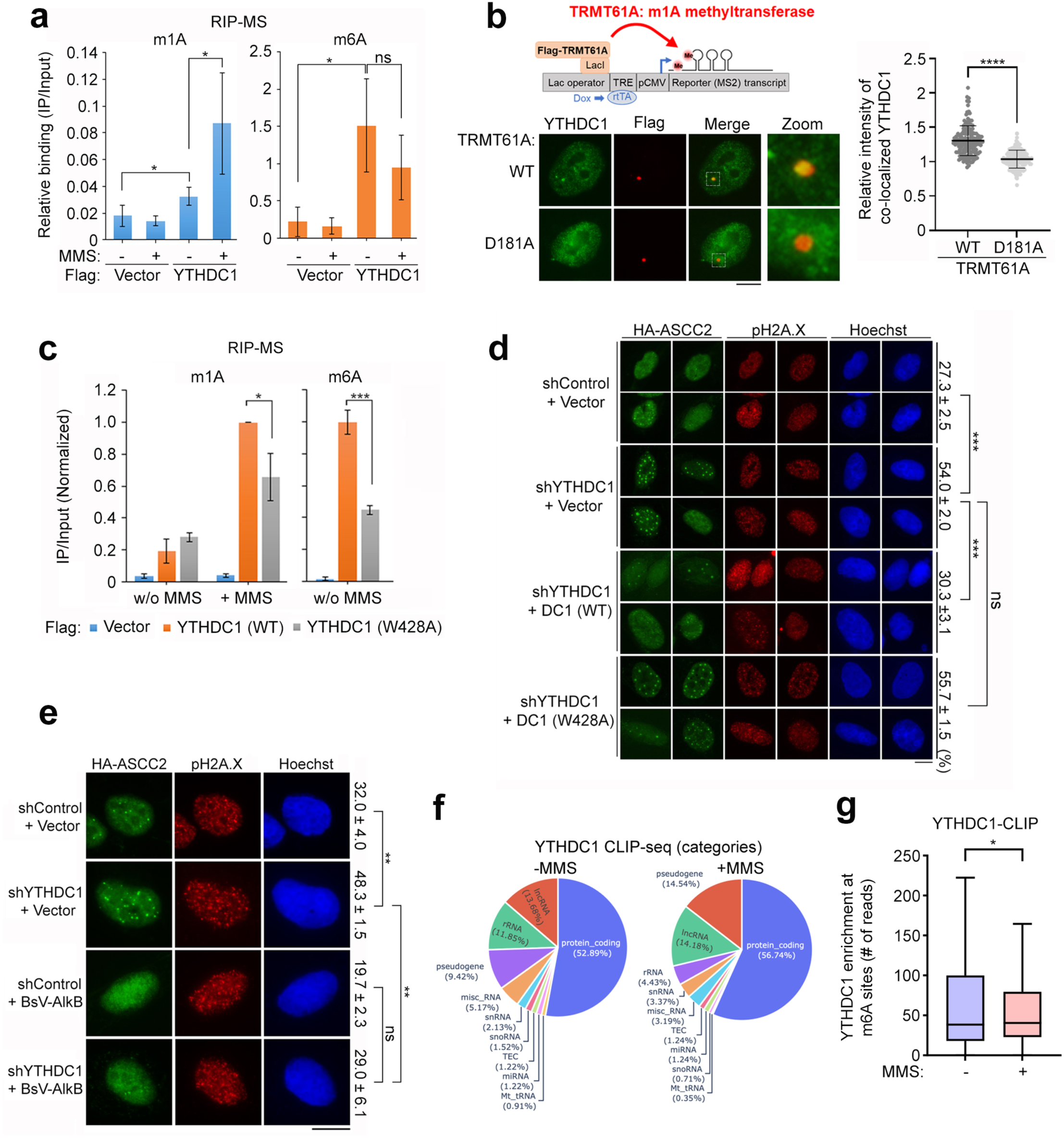
YTHDC1 binds to m1A-containing RNAs upon alkylation damage. **a**, 293T cells expressing Flag vector or Flag-YTHDC1 were harvested for RIP. Purified RNAs were digested, dephosphorylated, and the methylated nucleosides were analyzed by LC-MS/MS. The result is shown as the mean ± SD (n = 3 independent experiments; *p < 0.05 by Student’s t test; ns, non-significant). **b**, DD degron and Flag-tagged wildtype (WT) and catalytically inactive (D181A) TRMT61A-LacI expressing U2OS 2-6-3 single locus reporter cells were incubated with Shield1 ligand (300 nM) and doxycycline (1 µg/mL) for 24 h, then processed for immunofluorescence staining with the indicated antibodies. The representative images are shown below the schematic. Scale bar, 10 µm. Quantitation of YTHDC1 intensity is shown in the right panel. Error bars indicate the mean ± SD (n = 3 biological repeats, at least 50 cells were analyzed in each sample. ****p < 0.0001 by Mann-Whitney test). **c**, 293T cells expressing wildtype and YTH domain mutant (W428A) Flag-tagged YTHDC1 were harvested for RIP. Purified RNAs were digested, dephosphorylated, and the methylated nucleosides were analyzed by LC-MS/MS. The result is shown as the mean ± SD (n = 3 independent experiments; *p < 0.05, ***p < 0.001 by Student’s t test). **d**, **e**, HA-ASCC2 expressing U2OS cells were infected with the indicated lentiviral shRNAs and empty Flag vector and the indicated Flag-YTHDC1 (**d**), or Flag-BsV-AlkB (**e**) expression vectors. After MMS treatment, cells were processed for immunofluorescence staining with the indicated antibodies. Scale bar, 10 µm. % cells with more than 10 HA-ASCC2 foci are shown on the right of the representative images as the mean ± SD (n = 3 biological repeats, at least 100 cells were analyzed in each sample; **p < 0.01, ***p < 0.001 by Student’s t test; ns, non-significant). **f-g,** CLIP-seq analysis of YTHDC1 with (+) or without (-) MMS-induced damage. Categories of high confidence hits found in two repeats are shown in (**f**). Tukey boxplot of YTHDC1 enrichment at m6A sites from YTHDC1 CLIP-seq with or without MMS treatment (outliers not shown) is shown in (**g**). *p < 0.05 by using a nonparametric Wilcoxon matched-pairs signed rank test.

To further characterize the YTHDC1-m1A interaction, we performed an *in vitro* pulldown assay using GST-tagged YTH domain of YTHDC1 with m1A- or m6A-containing RNA oligonucleotides. GST-YTH^YTHDC1^ bound to the m1A as well as the m6A modification, suggesting a direct interaction between the YTH domain to m1A (Extended Data Fig. 3c). Furthermore, this interaction was reduced by addition of free m6A-containing RNAs (Extended Data Fig. 3c), which suggested a shared mode of recognition for m1A and m6A. To further probe the mechanism of binding, we used wildtype and mutant (W428A) YTH domain with either m1A or m6A RNAs (Extended Data Fig. 3d-e). Notably, this residue has been shown to be critical for the interaction between YTHDC1 and m6A^27^. Indeed, the GST-YTH^W428A^ mutant abrogated the binding to both m1A and m6A, further confirming that YTHDC1 may bind to m1A via its YTH domain (Extended Data Fig. 3d-e). Consistently, YTHDC1^W428A^ not only had reduced binding to methylated RNAs in cells, but also failed to rescue the ASCC foci formation phenotype, unlike wildtype YTHDC1 (Fig. 2c-d and Extended Data Fig. 3f-g). This supported a model where the recognition of methylated RNA is critical for the function of YTHDC1 in the alkylation response.

To distinguish the role of m1A versus m6A in this function of YTHDC1, we utilized the blueberry scorch virus AlkB protein (BsV-AlkB)^9, 28^, which we previously found to be specific for demethylating RNA and not DNA^9^. Further characterization of this enzyme demonstrated it to be highly specific for demethylating m1A and not m6A in RNA (Extended Data Fig. 4a-b). To determine whether this YTHDC1-dependent alkylation response on ASCC foci formation is mediated by m1A, we expressed an NLS-tagged BsV-AlkB in control and YTHDC1-depleted cells. Indeed, BsV-AlkB decreased ASCC foci formation in both control and YTHDC1-depleted cells to comparable levels (Fig. 2e and Extended Data Fig. 4c), which demonstrated that YTHDC1 regulates the ASCC pathway via recognizing m1A-containing RNAs.

Next, to identify RNA populations bound by YTHDC1 upon alkylation damage, we performed crosslinking immunoprecipitation followed by deep sequencing (CLIP-seq) with Flag-YTHDC1. The result showed that in both control and MMS-treated cells, YTHDC1 primarily interacted with protein-coding pre-mRNA regions, with more than 75% of these interactions occurring in introns (Fig. 2f and Extended Data Fig. 4d), consistent with its function in pre-mRNA processing^15–19^. Using this data, we determined the enrichment of the YTHDC1 CLIP-seq reads on m6A sites identified from a published database^29^. This showed that the overall YTHDC1 reads coverage on m6A sites is reduced in MMS-treated cells relative to untreated cells, indicating that the binding specificity of YTHDC1 to methylated RNAs may be altered upon alkylation damage (Fig. 2g and Extended Data Fig. 4e).

### YTHDC1 protects against RNA damage-induced DNA breaks

Since YTHDC1 loss increased alkylation sensitivity, we determined if this is due to the DNA damage response. Using the neutral comet assay^30^, we found that YTHDC1-depleted cells accumulated more DNA double-stranded breaks (DSBs) than control cells after alkylation damage (Fig. 3a-b). Strikingly, this phenotype was fully rescued by expressing the wildtype but not catalytically inactive NLS-tagged BsV-AlkB RNA demethylase (Fig. 3c and Extended Data Fig. 5a). These results indicated that in the absence of YTHDC1, chemically methylated RNAs may induce DNA breaks. Consistently, BsV-AlkB also rescued the alkylation hypersensitivity caused by YTHDC1 loss in a manner that depended on its catalytic activity (Fig. 3d).

**Fig. 3.**
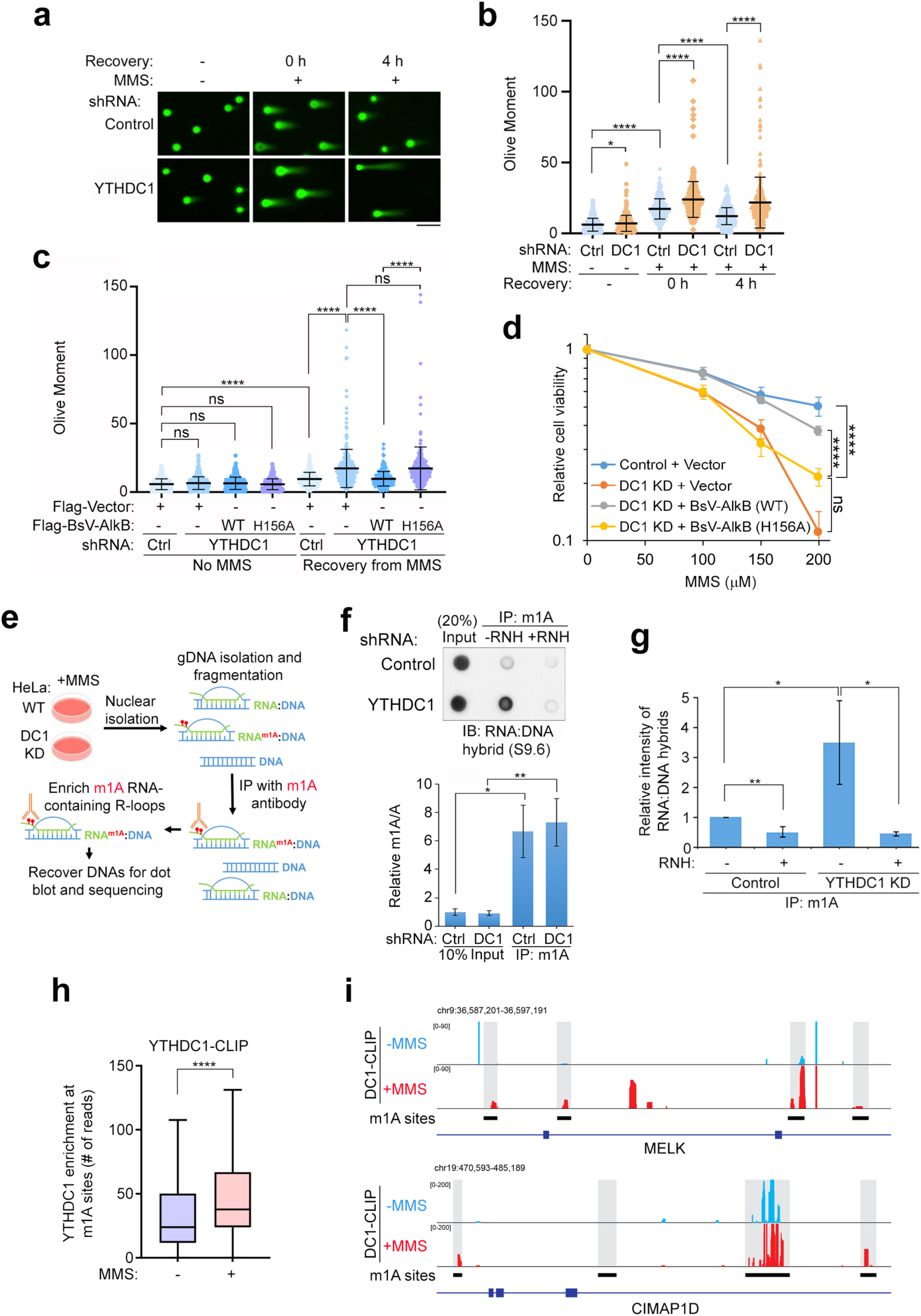
YTHDC1 protects against RNA damage induced DNA breaks. **a, b**, Neutral comet assay of control and YTHDC1 shRNA-expressing HeLa cells with or without MMS treatment. Representative images are shown in (**a**). Scale bar, 50 µm. The quantitation is shown in (**b**) (n = 3 biological replicates. At least 100 cells were analyzed in each sample. Error bars indicate mean the mean ± SD. *p < 0.05, **** p< 0.0001 by Mann-Whitney test). **c**, Neutral comet assay of control and YTHDC1 shRNA-expressing HeLa cells expressing empty Flag vector or the indicated Flag-BsV-AlkB expression vectors with or without MMS treatment. The quantitation is shown as the mean ± SD (n = 3 biological repeats. At least 100 cells were analyzed in each sample. **** p< 0.0001 by Mann-Whitney test, ns, non-significant). **d**, HeLa cells expressing inducible YTHDC1 shRNA were transduced with empty Flag vector or the indicated Flag-BsV-AlkB expression vectors with or without the induction of YTHDC1 shRNA with 0.5 µg/mL of doxycycline, then processed for MTS assay after treatment with the indicated concentration of MMS (n = 5 technical repeats for each sample). ****p < 0.0001 by two-way ANOVA test; ns, non-significant. **e**, Scheme of the m1A-DIP technique. After MMS treatment and isolation of nuclei, genomic DNA was purified followed by immunoprecipitation with m1A antibody. **f, g**, Dot blot of m1A-DIP samples probed with S9.6 antibody, with a representative blot shown in the upper panel of (**f**). The LC-MS/MS-based quantitation of m1A and A from genomic DNAs was shown in the bottom panel of (**f**) as the mean ± SD (n = 3 biological repeats; *p < 0.05, ** p< 0.01 by Student’s t test). The quantitation of three independent dot blot experiments is shown in (**g**) as the mean ± SD (*p < 0.05, ** p< 0.01 by Student’s t test). The intensity of S9.6 in each IP sample was normalized to the corresponding input, and depicted as relative to the control RNH (-) sample. **h,** Tukey boxplot of YTHDC1 enrichment at m1A sites from YTHDC1 CLIP-seq with or without MMS treatment (outliers not shown). ****p < 0.0001 by using a nonparametric Wilcoxon matched-pairs signed rank test. **i**, IGV tracks display for the m1A sites (shown in grey background) read coverage of YTHDC1 CLIP-seq reads in MELK and CIMAP1D genes.

Next, we investigated how damaged RNAs may induce DNA breaks. We previously showed that alkylation of a pre-mRNA can inhibit its splicing *in vitro*^9^; furthermore, inhibition of RNA processing is known to increase R-loops, which are three-stranded nucleic acid structures composed of an RNA:DNA hybrid and the associated non-template single-stranded DNA^31–33^. Previous proteomic studies have also suggested that YTHDC1 may associate with R-loops^34^; we confirmed this by immunoprecipitation of RNA:DNA hybrids using a monoclonal antibody (S9.6)^35^, which demonstrated an association with YTHDC1 in both normal and alkylation damage conditions (Extended Data Fig. 6a-b). Thus, the association of YTHDC1 with damaged RNAs may be in the context of R-loops. To test this, we first performed m1A DNA immunoprecipitation (m1A-DIP) using genomic DNA (gDNA) samples prepared from MMS-treated control and YTHDC1-depleted cells using an m1A-specific antibody (Fig. 3e and Extended Data Fig. 5b-c). Notably, this antibody enriched for m1A-containing but not m6A-containing RNAs in association with the genomic DNA (Extended Fig. 5c). To quantify m1A-associated R-loops, we used dot blotting to probe the m1A-DIP samples with the S9.6 antibody. This showed that in the absence of YTHDC1, the levels of m1A-containing R-loops was significantly increased (Fig 3f-g. and Extended Fig. 5c). We identified the genomic sites which contain m1A from the m1A-DIP samples and quantified the enrichment of YTHDC1 at these m1A sites through YTHDC1 CLIP-seq analysis. The results showed a significantly increased accumulation of YTHDC1 at these m1A sites upon MMS treatment, suggesting that alkylation damage induces the recruitment of YTHDC1 to the m1A sites (Fig. 3h-i and Extended Data Fig. 5d). Since these m1A sites were not homogeneous across the genome, we reasoned that this may reflect RNA accessibility. Indeed, we found that these m1A sites have higher RNA accessibility as shown by the higher NAI-N3 reactivity from icSHAPE (*in vivo* click selective 2-hydroxyl acylation and profiling experiment) analysis^36^ (Extended Data Figure 5e).

### YTHDC1 suppresses RNA damage-induced R-loops to promote genome integrity

To test whether YTHDC1 prevents RNA damage-induced R-loops, we performed R-loop quantitation using a recombinant GFP-tagged catalytically inactive RNase H1 as a probe (GFP-dRNH), which has been shown to bind with higher specificity to RNA:DNA hybrids compared to the S9.6 antibody in immunofluorescence studies^37^. We found that while YTHDC1 loss alone did not increase nuclear R-loops, these cells had significantly greater R-loop accumulation upon alkylation damage (Fig. 4a-b). Importantly, the GFP-dRNH signals were reduced by pre-treating the cells with a catalytically active RNaseH1^38^, indicating these signals are specific to RNA:DNA hybrids (Fig. 4a-b). To determine whether this accumulation of R-loops requires transcription, we pre-treated cells with the RNA polymerase II elongation inhibitor 5,6-dichloro-1-beta-D-ribofuranosylbenzimidazole (DRB)^39^. DRB reduced R-loop accumulation in YTHDC1-depleted cells to the same level as control cells (Fig. 4c). In addition, we found that this R-loop phenotype is associated with the role of YTHDC1 binding to damaged RNA, since the R-loop accumulation was rescued either by YTHDC1 in a YTH domain-dependent manner, or by BsV-AlkB in a catalytic activity-dependent manner (Fig. 4d-e and Extended Data Fig. 6c, respectively). These results suggested that both m1A demethylation and the YTH domain of YTHDC1 are required for repressing alkylation-induced R-loops.

**Fig. 4.**
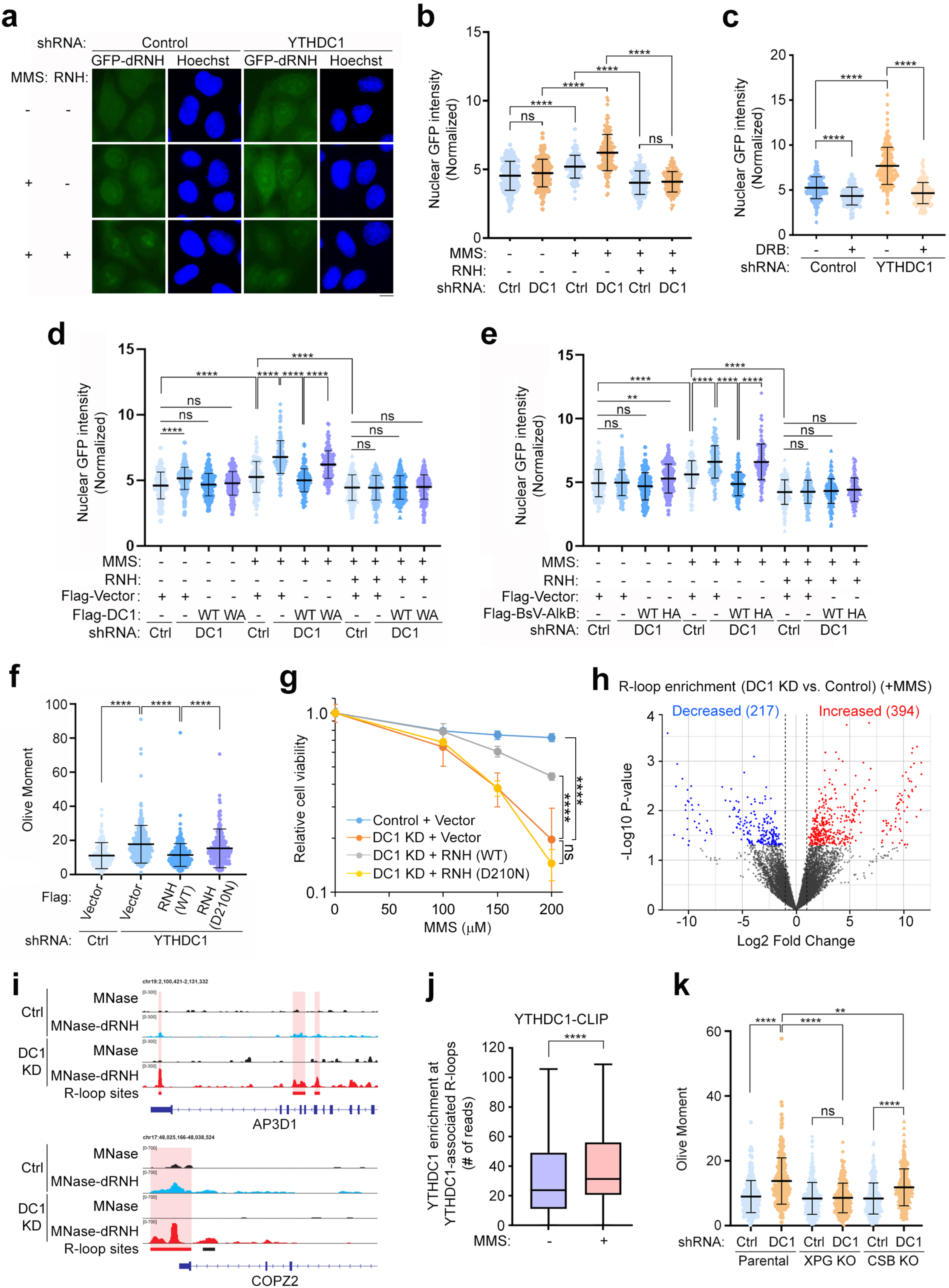
YTHDC1 suppresses RNA damage-induced R-loop formation to promote genome integrity. **a, b** HeLa cells expressing control and YTHDC1 shRNAs with or without MMS treatment were subjected to R-loop staining using purified GFP-dRNH. Representative images are shown in (**a**). Scale bar, 10 µm. The nuclear GFP intensity was quantified and shown in (**b**) as the mean ± SD (n = 3 biological repeats. At least 50 cells were analyzed in each sample. **** p< 0.0001 by Mann-Whitney test, ns, non-significant). **c**, HeLa cells were pre-treated with DMSO (-) or 50 µM DRB for 2 h, followed by MMS treatment. Cells were processed for R-loop staining using purified GFP-dRNH. Nuclear GFP intensity was quantitated and represented as the mean ± SD (n = 3 biological repeats. At least 50 cells were analyzed in each sample. **** p< 0.0001 by Mann-Whitney test; ns, non-significant). **d, e,** Control and YTHDC1 shRNA-expressing HeLa cells were transduced with empty Flag vector or the indicated Flag-YTHDC1 expression vectors (**d**), or the indicated Flag-BsV-AlkB expression vectors (**e**). Cells were treated with MMS as indicated, and then processed for R-loop staining using GFP-dRNH. Nuclear GFP intensity was quantitated and present by the mean ± SD (n = 3 biological repeats. At least 50 cells were analyzed in each sample. ** p < 0.01, **** p< 0.0001 by Mann-Whitney test; ns, non-significant). **f**, Neutral comet assay of control and YTHDC1 shRNA-expressing HeLa cells rescued with empty Flag vector or the indicated Flag-RNH expression vectors four hours after recovering from MMS exposure. The quantitation is shown as the mean ± SD (n = 3 biological repeats. At least 100 cells were analyzed in each sample. ****p < 0.0001 by Mann-Whitney test). **g**, HeLa cells expressing inducible YTHDC1 shRNA were transduced with empty Flag vector or the indicated Flag-RNH expression vectors with or without the induction of YTHDC1 shRNA with 0.5 µg/mL of doxycycline. After exposure to the indicated doses of MMS, cell viability was assessed with MTS assay (n = 5 technical repeats for each sample). ****p < 0.0001 by two-way ANOVA test; ns, non-significant. **h**, Volcano plot of MapR peaks identified in MMS-treated HeLa cells expressing control and YTHDC1 shRNAs. R-loop enrichment by YTHDC1 KD was quantitated and the significant (p < 0.05) peaks with the fold change Log2 > 1 (Increased) or Log2 <-1 (Decreased) were highlighted. **i**, IGV tracks display for the MapR read count coverage of the R-loop sites with increased enrichment in the YTHDC1 knockdown sample (shown in red background) in AP3D1 and COPZ2 genes. **j**, Tukey boxplot of YTHDC1 enrichment at YTHDC1-associated R-loops from YTHDC1 CLIP-seq with or without MMS treatment (outliers not shown). ****p < 0.0001 of MMS-treated samples compared to non-treated samples using a nonparametric Wilcoxon matched-pairs signed rank test. **k**, Neutral comet assay of wildtype, XPG KO, and CSB KO RPE-1 cells four hours after recovering from MMS exposure. The quantitation is shown as the mean ± SD (n = 3 biological repeats. At least 100 cells were analyzed in each sample. **p < 0.01, ****p < 0.0001 by Mann-Whitney test, ns, non-significant).

To further confirm that the R-loop accumulation upon YTHDC1 loss reflected a role of YTHDC1 in alkylation damage responses, we quantified DNA DSBs and cell viability in YTHDC1-depleted cells expressing wildtype and catalytically inactive (D210N) RNH. The expression of wildtype RNH significantly rescued the DNA DSBs and partially rescued the alkylation damage sensitivity induced by YTHDC1 depletion. However, the catalytically inactive RNH was not able to rescue either phenotype (Fig. 4f-g and Extended Data Fig. 6d). Therefore, YTHDC1 functions to suppress R-loop accumulation upon alkylation, impacting DNA DSB formation and alkylation damage sensitivity. We next performed MapR to identify R-loops genome-wide using a catalytically inactive RNH fused to micrococcal nuclease^40^. Moreover, we found that YTHDC1 depletion following MMS treatment resulted in increased enrichment of R-loops across the genome (Fig. 4h-i). In addition, the recruitment of YTHDC1 to these R-loops appeared to be alkylation-dependent, as higher YTHDC1 enrichment at these R-loops were observed upon MMS treatment (Fig. 4j and Extended Data Fig. 6e). Thus, YTHDC1 may play a role in mitigating the formation of R-loops induced by alkylation.

How do these alkylation-induced R-loops cause DNA DSBs? Unresolved R-loops may be processed by factors involved in transcription-coupled nucleotide excision repair (TC-NER), which in turn generate DNA DSBs^41^. We tested this possibility by using the parental and TC-NER-deficient (XPG and CSB knockout) RPE-1 cells and performed comet assays after recovery from MMS exposure. The result showed that cells deficient for the XPG nuclease rescued DNA DSB formation upon YTHDC1 loss, while CSB knockout cells partially rescued these DSBs (Fig. 4k). This demonstrated that these alkylation-induced R-loops in YTHDC1-depleted cells are likely processed by nucleases associated with TC-NER.

### Induction of m1A is sufficient to induce DNA breaks in YTHDC1 or THOC depleted cells

The fact that BsV-AlkB was able to rescue DNA DSBs induced by alkylation in the YTHDC1-depleted cells suggested that m1A in RNA is necessary for induction of DNA breaks (Fig. 3c). We next tested whether induction of nuclear m1A is sufficient to induce DNA DSBs upon YTHDC1 loss. To this end, we expressed an NLS-tagged m1A methyltransferase complex, TRMT6/61A^42^ and confirmed the nuclear expression of individual TRMT components (Extended Data Fig. 7a). We found that expressing TRMT6/61A in the nucleus did not induce DSBs in control cells. However, this RNA methyltransferase was sufficient to induce DNA breaks in the YTHDC1-depleted cells in a manner dependent on TRMT61A catalytic activity (Fig. 5a-c and Extended Data Fig. 7b-c). These results suggested that induction of RNA damage is sufficient to generate DSBs in the YTHDC1-depeted cells. Furthermore, by using the single locus reporter system, we demonstrated that targeting of this RNA methyltransferase induced phosphorylated ATM (pATM), a well-established DNA DSB marker^43^, at the locus site in the YTHDC1-depleted cells (Fig. 5d-e). Thus, m1A-induced DSBs may occur upon YTHDC1 loss, and these breaks are likely in *cis* relative to the site of RNA damage. THOC2 depletion also resulted in greater DSB formation upon expression of the nucleus-localized TRMT6/61A (Fig. 5f-g and Extended Data Fig. 7d), consistent with the notion that YTHDC1 may function with THOC to suppress damaged RNA-induced DNA DSBs. We term this phenomenon <u>R</u>NA <u>d</u>amage-induced DNA <u>b</u>reaks (RDIBs).

**Fig. 5.**
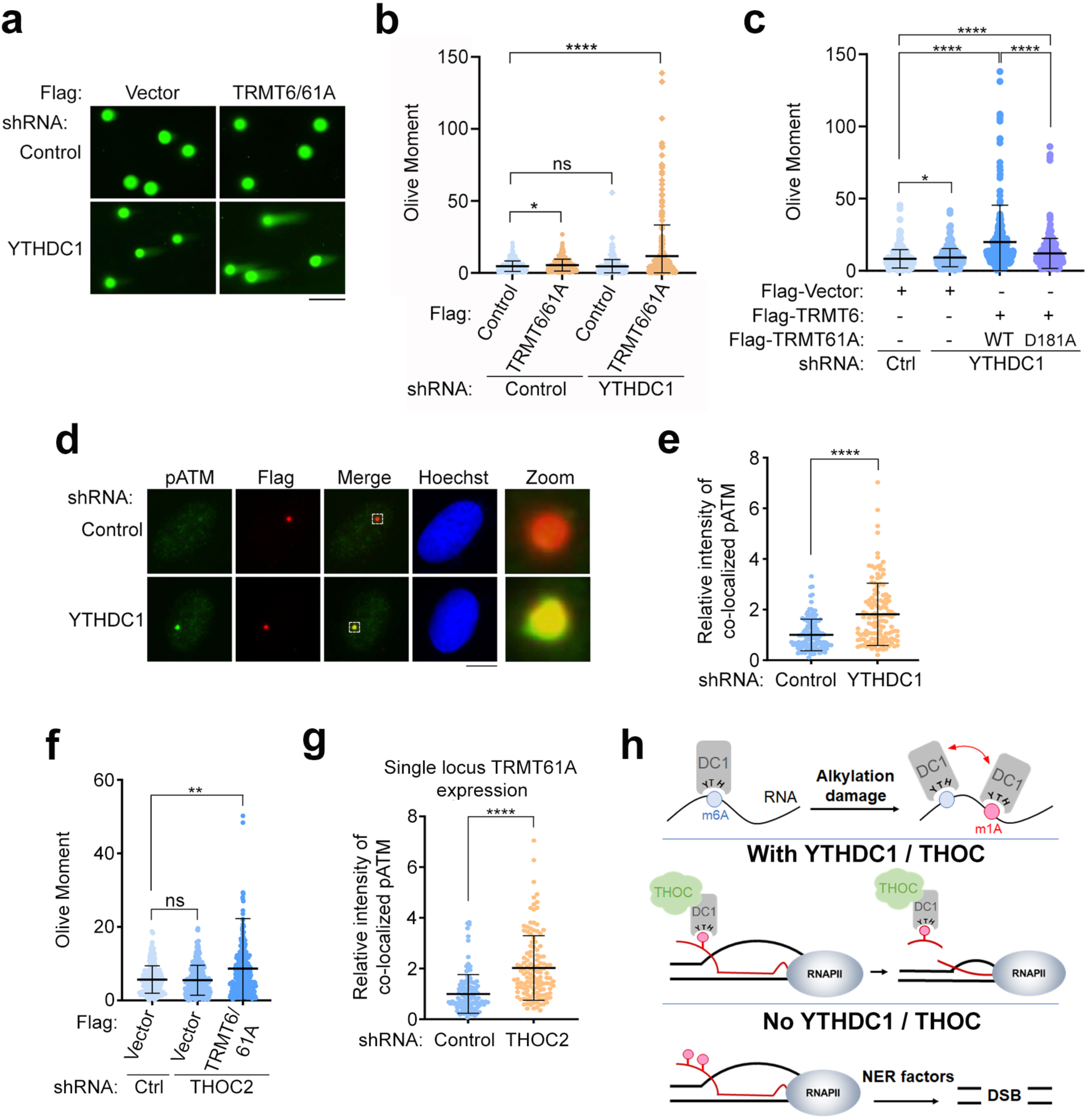
Induction of m1A is sufficient to generate DNA breaks in the absence of YTHDC1 or THO complex. **a, b,** Control and YTHDC1 shRNA-expressing HeLa cells were transduced with empty Flag vector or the NLS fused Flag-TRMT expression vectors for 48 hours. Double-stranded breaks were assessed using neutral comet assay. The representative images are shown in (**a**). Scale bar, 50 µm. The quantitation is shown in (**b**) as the mean ± SD (n = 3 biological repeats. At least 100 cells were analyzed in each sample. *p < 0.05, ****p < 0.0001 by Mann-Whitney test; ns, non-significant). **c**, Control and YTHDC1 shRNA-expressing HeLa cells expressing empty Flag vector and the indicated NLS fused Flag-TRMT expression vectors. Neutral comet assay was performed as in **(b)**. The quantitation result is shown as the mean ± SD (n = 3 biological repeats. At least 100 cells were analyzed in each sample. *p < 0.05, ****p < 0.0001 by Mann-Whitney test). **d-e,** The DD degron/Flag-tagged TRMT61A-LacI expressing U2OS 2-6-3 single locus reporter cells were infected with lentiviral control or YTHDC1 shRNAs. After 48 hours, cells were incubated with 300 nM Shield1 ligand and 1 µg/mL doxycycline for 24 hours, then processed for immunofluorescence staining using the indicated antibodies. The representative images are shown in (**d**). Scale bar, 10 µm. Quantitation of pATM staining is shown in (**e**). Error bars indicate the mean ± SD (n = 6 biological repeats, more than 20 cells were analyzed in each sample. ****p < 0.0001 by Mann-Whitney test). **f**, Control and THOC2 shRNA-expressing HeLa cells were transduced with empty Flag vector or Flag-NLS-TRMT expression vectors. Double-stranded breaks were assessed using neutral comet assay. The quantitation result is shown as the mean ± SD (three independent experiments. At least 100 cells were analyzed in each sample. **p < 0.01 by Mann-Whitney test; ns, non-significant). **g**, The DD degron/Flag-tagged TRMT61A-LacI expressing U2OS 2-6-3 single locus reporter cells were infected with lentiviral control or THOC2 shRNAs. After 48 hours, cells were incubated with 300 nM Shield1 ligand and 1 µg/mL doxycycline for 24 h, then processed for immunofluorescence staining using the indicated antibodies. Quantitation result is shown as the mean ± SD (n = 3 biological repeats, more than 40 cells were analyzed in each sample. ****p < 0.0001 by Mann-Whitney test). **h,** The proposed model: (Top) YTHDC1 binds to m1A-containing RNAs during alkylation damage. (Middle) YTHDC1 works in conjunction with the THO complex, helping to remove damaged nascent RNAs. (Bottom) In the absence of YTHDC1 or THOC, cells accumulate R-loops containing due to aberrantly methylated RNAs, resulting in DNA double-stranded breaks (DSBs) through processing by TC-NER factors.

In sum, our work has revealed two major findings: First, the YTHDC1 and THOC are important for alkylation responses in cells in a manner that depends on the recognition of an aberrant RNA methylation mark induced by chemical agents. Second, in the absence of YTHDC1 or THOC, these aberrantly methylated RNAs induce R-loops, which are then processed into DNA DSBs via TC-NER factors (Fig. 5h). These data provide the first definitive evidence that damaged RNAs impact genome integrity in the context of alkylation stress and identify a complex that prevents their accrual. Further study will be necessary to better determine how YTHDC1 and THOC resolve damaged RNA-associating R-loops to promote genome stability. Besides YTHDC1, recent reports have also found that HRSP12 and TDP-43 bind to m1A-containing RNAs to mediate methylated RNA-mediated functions^44, 45^. Whether these functions are perturbed by alkylation damage, which induces m1A, will be important to elucidate.

## Materials and Methods

### Cell culture

All human cell lines were cultured in Dulbecco’s modified eagle medium (Invitrogen), supplemented with 10% fetal bovine serum (Atlanta Biologicals), 100 U/ml of penicillin-streptomycin (Gibco) at 37°C and 5% CO_2_. Cell lines was authenticated by using the ATCC human STR profiling service and regularly tested for mycoplasma contamination by using PCR-based Mycoplasma Testing at the Washington University Genome Engineering and iPSC Center (GEiC). Any genetic manipulations by CRISPR/Cas9 were verified by DNA sequencing at the GEiC at Washington University. p53-deficient RPE-1 (parental, XPG KO, and CSB KO) cells were kindly provided by Dr. Martijn Luijsterburg (Leiden University Medical Center) and knockouts were verified by deep sequencing at the GEiC at Washington University. The U2OS 2-6-3 reporter cell line^9, 23^ is maintained in the culture media with hygromycin (100 µg/mL) and blasticidin (5 µg/mL). Preparation of viruses, transfection, and viral transduction were performed as described previously^6^. Knockdown experiments (using lentiviral shRNA) were performed by infecting cells with the indicated lentivirus and selecting with puromycin (1 µg/mL) for 48-72 hours. For the doxycycline-inducible knockdown, cells were incubated with 0.5 µg/mL doxycycline (Sigma) for at least 72 hours.

### Plasmids

For mammalian cell expression, YTHDC1, RNaseH1, TRMT cDNAs or gBlocks were subcloned into pHAGE-CMV-Flag vector by Gateway recombination as described^6^. The pHAGE-3xHA-ASCC2, pMSCV-Flag-RNF113A, pHAGE-Flag-BsV-AlkB-NLS (WT and H156A), pHAGE-Flag-TRMT61A-NLS (WT and D181A) vectors was cloned previously^6, 9^. For the degron tagged expression, WT and D181A Flag-TRMT61A-NLS are cloned into the pLVS-PTuner-mCherry-LacI vector which was generated as previously described^9^. For recombinant protein purification, WT and W428A YTH domain of YTHDC1 (amino acids 337-509) were cloned into pGEX-4T1 using BamH1/XhoI restriction enzymes^13^. All the constructs derived by PCR or from gBlocks were confirmed by Sanger sequencing. For shRNA-mediated knockdown, all TRC pLKO.1 vectors (targeting the YTH domain-containing proteins, THOC1, and THOC2) were purchased from Sigma. The TRIPZ doxycycline (Dox)-inducible lentiviral YTHDC1 shRNA vector is purchased from Dharmacon.

### Immunofluorescence microscopy

All immunofluorescence microscopy for foci analysis was performed as previously described^6^, with minor modifications. For the MMS-induced ASCC foci, U2OS cells were treated with 500 µM MMS in culture medium at 37°C for six hours, washed with 1× PBS, then extracted with 1× PBS containing 0.2% Triton X-100 and protease inhibitors for 20 minutes on ice. Cells were fixed with 3.2% paraformaldehyde for 20 minutes at room temperature. For the single locus recruitment, the U2OS 2-6-3 cells expressing degron-tagged LacI fusion proteins were incubated with 300 nM Shield1 ligand (Takara Bio) and 1 µg/mL doxycycline for 24 hours, then washed with 1× PBS prior to extraction and fixation as above. For the NLS-tagged TRMT staining, cells were directly fixed with paraformaldehyde without Triton X-100 extraction. All cells were washed extensively with IF Wash Buffer (1× PBS, 0.5% NP-40, and 0.02% NaN_3_), then blocked with IF Blocking Buffer (IF Wash Buffer plus 10% FBS) for at least 30 minutes. Primary antibodies were diluted in IF Blocking Buffer overnight at 4°C. After staining with secondary antibodies (conjugated with either Alexa Fluor 488 or 594; Millipore) and Hoechst 33342 (BD Biosciences), samples were mounted using Prolong Gold mounting medium (Invitrogen). Epifluorescence microscopy was performed on an Olympus fluorescence microscope (BX-53) using an UPlanS-Apo 100X/1.4 oil immersion lens and CellSens Dimension software. Raw images were exported into Adobe Photoshop, and for any adjustments in image contrast or brightness, the levels function was applied. For foci quantitation, at least 100 cells were analyzed in triplicate, unless otherwise indicated. The fluorescence intensities were quantitated by ImageJ. For the single locus recruitment, the total intensity of the Flag-TRMT61A-LacI co-localized YTHDC1 or pATM was normalized with its adjacent nuclear region as background.

### Cell survival assay

For DNA damaging agent viability assay, HeLa (2500 cells/well) or U2OS (3000 cells/well) were plated and cultured overnight in 96-well plates. Cells were then exposed to medium containing the indicated concentration of MMS (Millipore Sigma) for 24 hours at 37 °C. The media was then replaced with normal media, and cell viability was performed using the MTS assay (Promega) 72 hours after initial damaging agent exposure. All the experiments were carried out in technical quintuplicate with at least two biological repeats. For the colony formation assay, YTHDC1 KD or control HeLa cells were trypsinized, counted, and plated at low density. After overnight incubation, the cells were treated with the indicated doses of MMS for 24 hours in complete media. MMS-containing media was replaced with fresh media and cells were incubated for 14 days, fixed, and stained with crystal violet. The experiment was performed in triplicate for each cell line and drug dose. Colonies were counted and relative survival was normalized to untreated controls.

### Endogenous knock-in by CRISPR-Cas9 system

The small guide RNA (sgRNA) for targeting endogenous YTHDC1 and single-stranded donor DNA containing a Flag-tag sequence are designed by using the online Alt-R™ CRISPR HDR Design Tool from Integrated DNA Technology (IDT). Cas9 protein was purchased from Millipore Sigma (A36498). Transfection of sgRNA, donor DNA and Cas9 protein into HeLa-S cells was conducted by using Neon Transfection Electroporation System following manufacturer’s manual (Thermo Fisher Scientific). The expression of the endogenously Flag tagged YTHDC1 in the pooled cells was determined by western blotting three days after electroporation. A positive single colony was chosen up and expanded, followed by validation using western blotting and deep sequencing.

### YTHDC1 complex purification and proteomic analysis

Parental and endogenously Flag-tagged YTHDC1-expressing HeLa-S cells with or without MMS treatment were harvested for nuclear extraction according to previous protocols^6^. The YTHDC1 complex was purified using anti-Flag (M2) resin (Millipore Sigma) in TAP buffer (50 mM Tris-HCl pH 7.9, 100 mM KCl, 5 mM MgCl_2_, 10% glycerol, 0.1% NP-40, 1 mM DTT, and protease inhibitors). After peptide elution, the complexes were TCA precipitated and associated proteins were identified by LC-MS/MS at the Taplin Mass Spectrometry Facility (Harvard Medical School) using an LTQ Orbitrap Velos Pro ion-trap mass spectrometer (ThermoFisher) and Sequest software^46^. Total and unique peptide spectral counts, as well as total peptide intensities, can be found in Extended Data Table 1.

### Immunoprecipitation

Immunoprecipitation of Flag-YTHDC1 was performed by using parental and the endogenously Flag-tagged YTHDC1-expressing HeLa-S cells. The cells were resuspended in high-salt buffer (50 mM Tris-HCl pH 7.9, 300 mM KCl, 10% glycerol, 1.0% Triton X-100, 1 mM DTT, and protease inhibitors), lysed by sonication, and centrifuged. An equal volume of buffer without KCl was added, and the lysate was incubated with anti-Flag (M2) resin (Millipore Sigma). After incubation at 4°C with rotation, the resins were washed extensively with buffer containing 150 mM KCl. Bound material was eluted with Laemmli buffer and analyzed by Western blotting. For immunoprecipitation of RNA:DNA hybrids, nuclear extraction of cells were performed by incubating with the extraction buffer (85 mM KCl, 5 mM PIPES pH8.0, 0.5% NP-40, and protease inhibitors) on ice for 10 minutes prior to lysis and sonication in high-salt buffer. The lysate was incubated with S9.6 or normal mouse IgG overnight at 4°C and the immunocomplex were pulled down by protein G agarose (Pierce). After washing extensively, bound material was eluted with Laemmli buffer and analyzed by Western blotting.

### RNA immunoprecipitation coupled with nucleoside mass spectrometry (RIP-MS)

The RIP-MS was performed as described^9, 20^. HeLa-S or 293T cells expressing empty pMSCV-Flag or pHAGE-Flag vector, pMSCV-Flag-RNF113A, pHAGE-Flag-YTHDC1 (WT) or YTH domain mutation (W428A) grown in 15-cm dishes were treated with or without 5 mM MMS for 1 hour. Cells were harvested and the pellets were lysed with 2 volumes of lysis buffer (10 mM HEPES pH 7.5, 150 mM KCl, 0.5% NP-40, 2mM EDTA, 10 mM β-ME, protease inhibitors and 400 U/ml RNase inhibitor), incubated on ice for 5 minutes then shock-frozen at −80°C for 30 minutes. Cell lysates were thawed on ice and centrifuged at 15,000g for 15 minutes. The supernatant was immunoprecipitated with Anti-Flag (M2) resin at 4°C for 4 hours. Beads were washed eight times with 1 ml ice-cold NT2 buffer (50 mM HEPES pH 7.5, 200 mM NaCl, 0.05% NP-40, 2mM EDTA, 10 mM β-ME and 200 U/ml RNase inhibitor) and eluted with 5 packed bead volumes of NT2 buffer containing 0.4 mg/ml Flag peptide. The supernatant was mixed with 1 ml TRIzol for RNA extraction according to the manufacture’s protocol, then digested for nucleoside mass spectrometry analysis.

### Nucleoside Mass Spectrometry

Samples were digested to nucleosides at 37°C overnight with Nuclease S1 from *Aspergillus oryzae* (Millipore Sigma), followed by dephosphorylation with FastAP alkaline phosphatase (Thermo Fisher Scientific) at 37°C for 1 hour. Chromatographic separation was performed using an Agilent 1290 Infinity II UHPLC system with a ZORBAX RRHD Eclipse Plus C18 2.1 x 50 mm (1.8 um) column. The mobile phase consisted of water and methanol (with 0.1% formic acid) run at 0.5 ml/min. The run started with a 3 min gradient of 2-8% methanol, followed by a sharp increase to 98% methanol which was maintained for 4 min. Mass spectrometric detection was performed using an Agilent 6470 Triple Quadrupole system operation in positive electrospray ionization mode, monitoring the mass transitions 282/150 (for m1A and m6A), 268.1/136 (for A), 258/126 (for m3C), 244.1/112 (for C), 298/166 (for m7G), and 284.2/152 (for G). The signals of m1A and m6A is distinguished by the retention times described previously^20^.

### Tandem ubiquitin binding element (TUBE) assay

HeLa-S cells (∼6-8 x 10^7^) grown in a spinner flask were treated with or without 0.5 mM MMS for four hours at 37°C. The cells were collected by centrifugation and washed with ice-cold PBS and frozen at −80°C as two cell pellets. Each cell pellet was resuspended in 10 ml TUBE lysis buffer (50 mM Tris-HCl pH 7.5, 1 mM EGTA, 1 mM EDTA, 1% (v/v) Triton X-100, and 0.27 M sucrose) containing freshly added 100 mM iodoacetamide, protease and phosphatase inhibitors, then rotated at 4°C for one hour for lysis. The extract was spun at 6500 rpm for 30 minutes. Supernatant was transferred to a fresh tube and spun at 6500 rpm for 5 minutes. A fraction of the spun extract was kept as input, and the rest was rotated overnight at 4°C with 50 µL commercial ubiquitin-conjugated TUBE beads (Boston Biochem). The beads were washed three times with 10 ml high salt TAP buffer (50 mM Tris-HCl pH 7.9, 300 mM KCl, 5 mM MgCl_2_, 0.2 mM EDTA, 0.1% NP-40, 10% glycerol, 2 mM β-ME, 0.2 mM PMSF) and twice with 1 ml low salt TAP buffer (50 mM Tris-HCl pH 7.9, 0 mM KCl, 5 mM MgCl_2_, 0.2 mM EDTA, 0.1% NP-40, 10% glycerol, 2 mM β-ME, 0.2 mM PMSF). Beads were resuspended in 50 µL Laemmli buffer and analyzed by Western blotting.

### Protein purification

Recombinant His-tagged BsV-AlkB (WT and H156A) proteins were purified according to the previous descritptions^9^. Recombinant GST and GST-YTHDC1-YTH (WT and W428A) were purified as previously described^27^. Briefly, GST vectors were transformed into Rosetta (DE3) cells. When the culture reached OD 600 ∼ 0.6, the protein expression was induced by 1 mM isopropyl 1-thio-β-D-galactopyranoside (IPTG, Sigma) at 16°C overnight. The recombinant proteins were subsequently extracted with glutathione agarose (Pierce), following the manufacturer’s recommended procedures. Recombinant GFP-dRNH was purified by the Expression and Molecular Biology (EMB) Core of the Structural Cell Biology of DNA Repair Machines (SBDR) Program, by following the protocol as described previously^37^. The GST-MNase and GST-RNΔ-MNase proteins for MapR analysis were purified according to the previous description^40^.

### *In vitro* pulldown assays

For the GST pulldown, 0.2 µM of GST proteins are incubated with indicated concentrations of RNA oligos in 200 µL of TAP buffer (50 mM Tris-HCl pH 7.9, 100 mM KCl, 0.1% NP-40, 5 mM MgCl_2_, 0.2 mM EDTA, 10% glycerol, 0.2 mM PMSF, 10 mM β-ME) supplied with protease inhibitors and an RNase inhibitor (NEB, M0314L) for 1 hour at 4°C. The protein-RNA complexes were pulled down by pre-washed glutathione agarose for 1 hour at 4°C. Agarose beads were washed three times with high salt TAP buffer (50 mM Tris-HCl pH 7.9, 300 mM KCl, 0.1% NP-40, 5 mM MgCl_2_, 0.2 mM EDTA, 10% glycerol, 0.2 mM PMSF, 10 mM β-ME) and twice with TAP buffer supplemented with protease and RNase inhibitors. The agarose was mixed with 1 mL Trizol for RNA extraction according to the manufacture’s protocol, then digested for nucleoside mass spectrometry analysis. For the Streptavidin pulldown, 0.25 µM of biotin-labelled RNA oligos were incubated with streptavidin agarose resin (Pierce) in 200 µL of TAP buffer supplied with protease inhibitors and an RNase inhibitor for 1 hour at 4°C. After incubation, oligo-bound resin was washed three times with TAP buffer prior to incubated with 1 µg of GST and GST-fused proteins in 200 µL of TAP buffer for 1 hour at 4°C. The resin containing protein-oligo complexes were washed three times with high salt TAP buffer followed by twice with TAP buffer, and then suspended with Laemmli sample buffer prior Western blotting with the indicated antibody. The RNA oligo sequence is 5’-CAGAGGAGGUAAAAAAAUGGXCUUGUACAAA-3’ (X = A, m1A, or m6A).

### Nucleic acid demethylase assay

Methylated RNA oligonucleotides (sequences shown in above *In vitro* pulldown assays) were purchased from Midland Certified Reagents Company (Midland, TX) and TriLink Biotechnologies. Demethylation reactions were carried out using 60 pmol of substrate oligonucleotide in the presence of 20 pmol BsV-AlkB protein for 1 hour at 37°C in a 50 µl reaction mixture containing 50 mM HEPES–KOH pH 7.5, 2 mM ascorbic acid, 100 µM 2-oxoglutarate, 40 µM FeSO_4_ and 1 µl murine RNase inhibitor (NEB). The products of the reactions were digested to nucleosides and analyzed by LC-MS/MS, as described above (Nucleoside Mass Spectrometry).

### YTHDC1 crosslink immunoprecipitation (YTHDC1-CLIP)

Ten 15-cm dishes of HeLa cells expressing pHAGE-Flag-YTHDC1 were used for each CLIP. Cells were treated with or without 5 mM MMS for 1 hour and crosslinked by exposing twice with 0.15 J/cm^2^ of 254 nm UV, then scraped off and lysed with 3 volumes of 1X NP-40 buffer (50 mM HEPES-KOH pH7.5, 150 mM KCl, 2 mM EDTA, 0.5% NP-40, 10 mM β-ME, protease inhibitors and 400 U/ml RNase inhibitor). Clear cell lysates were treated with RNase T1 (Thermo Fisher Scientific) to a final concentration of 0.1 U/μl at room temperature for 15 minutes and immunoprecipitated with anti-Flag magnetic beads (Millipore Sigma) at 4°C for 1 hour. Beads were washed three times with IP-wash buffer (50 mM HEPES-KOH pH7.5, 300 mM KCl, 0.05% NP-40. 10 mM β-ME, protease inhibitors and 200 U/ml RNase inhibitor) and treated with RNase T1 to a final concentration of 10 U/μl at room temperature for 15 minutes. SUPERase•In™ RNase Inhibitor (Thermo Fisher Scientific) was then added to a final concentration of 1 U/μl to inhibit RNase T1. Beads were further washed three times with high-salt wash buffer (50 mM HEPES-KOH pH7.5, 500 mM KCl, 0.05% NP-40. 10 mM β-ME, protease inhibitors and 200 U/ml RNase inhibitor) and twice with PNK buffer (50 mM Tris-HCl pH7.5, 50 mM NaCl, 50 mM MgCl_2_). Beads were treated with 10 μl of T4 Polynucleotide kinase (PNK; NEB) in 100 μl of 1X T4 PNK Reaction buffer containing 1 U/μl SUPERase•In™ RNase Inhibitor at 37°C for 30 minutes, followed by addition of 1 mM ATP and 5 μl PNK for another 30 minutes incubation. After removal of the supernatants, beads were incubated with proteinase K at 55°C for 30 minutes with shaking. The supernatants were collected and RNAs were purified by using RNA Clean & Concentrator-5 kit (Zymo Research). The purified RNAs were subjected to library preparation using a SeqPlex RNA Amplification Kit (Millipore Sigma) and sequenced on an Illumina NovaSeq 6000 in the Genome Technology Access Center at Washington University.

### Neutral comet assay

HeLa or RPE-1 cells were plated 24 hours before treatment with 0.25 mM MMS for 1 hour. Media was replaced and incubated for another 4 hours when indicated. Cells were then trypsinized and resuspended to 2 X 10^5^ cells/mL in cold PBS. Cells were combined with low melting point agarose 1:10 and spread onto a comet slide (Trevigen) and allowed to dry at 4°C for 30 minutes. Slides were placed in lysis solution (Trevigen) at 4°C for 1 hour before immersion in 1X Electrophoresis buffer for 30 minutes. Lysed cells were then electrophoresed at 25V for 45 minutes at 4°C. Slides were incubated with DNA precipitate solution (1M ammonium acetate, 95% ethanol) for 30 minutes, and subsequently washed in 70% ethanol for 30 min. Slides were then dried overnight at room temp. Slides were stained with 1X SYBR Gold (Thermo Fisher Scientifgic) for 30 min, and images were acquired with a fluorescence microscope (Olympus BX-53) using a 10X lens and CellSens Dimension software. At least 100 cells were analyzed for each sample and the levels of DNA breaks was quantitated using the CometScore software.

### m1A and S9.6 DNA immunoprecipitation (m1A- and S9.6-DIP)

HeLa cells were treated with 2 mM MMS for 1 h when indicated. After nuclear extraction, genomic DNAs were purified by using the DNeasy Blood & Tissue Kit (Qiagen). The DNA was then fragmented to 100-600 bp by sonication. For the RNH digestion, gDNA was treated with RNase H (NEB, M0297S) (5 U per µg of gDNA) in 1× RNase H buffer overnight at 37 °C before immunoprecipitation and recovered using the Oligo Clean & Concentrator kit (Zymo Research). For m1A-DIP, gDNA was denatured at 95°C for 10 minutes right before immunoprecipitation. 10 µg of gDNA was used for immunoprecipitation with 2 µg of m1A, S9.6, or normal mouse IgG antibodies, and then the immunocomplex was pulled down by protein G magnetic beads (Thermo Fisher Scientific, 10003D). After extensively washing, beads were digested with protease K and the DNA was recovered using the Oligo Clean & Concentrator kit. For the dot blot assay, the purified DNA was dropped onto a positively charged nylon memberane (Millipore Sigma, 11209299001) and the membrane was crosslinked by exposing with 0.12 J/cm^2^ of 254 nm UV. After washing by 1X PBS, the membrane was blocked with 5% milk/TBST, then processed with immunoblotting with S9.6 antibody. For m1A and m6A quantitation, DNAs were digested overnight with the nucleoside digestion mix prior to nucleoside mass spectrometry. For m1A-DIP-seq, DNA samples were subjected to library preparation and sequenced by Azenta Life Science.

### RNA-DNA hybrid (R-loop) staining and quantitation by using GFP-dRNH

R-loop staining was performed by using GFP-dRNH according to the previous descriptions^36^. In brief, cells were treated with 1 mM MMS for 2 h to induce R-loop formation, then recovered from MMS exposure with fresh media for another 2 h. Cells were fixed with ice-cold methanol for 5 minutes at −20°C and then washed three times with 1XPBS. For the RNaseH treatment, cells were then incubated with RNase H (NEB, 1: 50 in 1X RNase H buffer) at 37°C for 4 hours. Cells were blocked with 3% BSA in 1X PBS for 30 minutes at room temperature and incubated with GFP-dRNH (0.23 µg/µL, diluted 1: 1000 in the Blocking buffer) and Hoechst at 37°C for 1.5 hours. After incubation, cells were washed three times with IF Wash Buffer (1× PBS, 0.5% NP-40, and 0.02% NaN_3_) and mounted for epifluorescence microscopy. Images were analyzed using ImageJ that sequentially detected and outlined nuclei based on Hoechst staining, then subtracted background intensity and measured mean intensity of GFP in nuclei to avoid bias. At least 50 cells were counted for each individual replicate.

### MapR

Control or YTHDC1 shRNA-expressing HeLa cells were treated with 2 mM MMS for 1 h, then subjected to MapR experiment by following the established procedures with minor changes^47^. Briefly, cells were trypsinized, suspended, and immobilized with concanavalin A-coated magnetic beads (Polyscience # 86057) by rotating at room temperature for 1 hour. After washing, cell-bound beads were resuspended with 50 µL dig-wash buffer (5% digitonin, 20 mM HEPES pH 7.5, 150 mM NaCl, 0.5 mM spermidine, protease inhibitors) and incubated overnight with 1 µM GST-RNΔ-MNase or GST-MNase by rotating at 4°C. Nucleic acids were then released from chromatin by activating MNase using CaCl_2_. After protease K treatment, DNAs were recovered using the MiniElute® PCR purification kit (Qiagene #28004). The purified DNAs were subjected to library preparation and sequenced on an Illumina NovaSeq X Plus in the Genome Technology Access Center at Washington University.

### Quantification of YTHDC1-CLIP reads coverage on m6A, m1A, and YTHDC1-associated R-loop sites

The sequencing reads from m6A IP were obtained from Yang *et al*. (2022)^29^ (GEO: SRR20018907, SRR20018904, SRR20018897, SRR20018896). The m1A-DIP, YTHDC1-CLIP, and MapR were initially processed by adapter trimming using TrimGalore (version 0.6.0). Subsequently, these reads were aligned to the GENCODE Release 45 (GRCh38.p14) genome. This alignment was executed using STAR (version 2.5.1a) for m6A IP and YTHDC1-CLIP, and Bowtie2 (version 2.4.1) for m1A IP and MapR. To determine m6A and m1A sites, we utilized the MACS2 (version 2.1.1) callpeak module with the *--keep-dup all* option. For each m1A and m6A IP sample, its corresponding input RNA sample served as the background control in peak calling. To determine R-loop sites, we utilized the MACS2 (version 2.1.1) callpeak module with the *--keep-dup all* option from the MapR GST-RNΔ-MNase treated sample while using the MapR GST-MNase treated sample as the background input control in peak calling. The final m6A and m1A sites were determined based on a criterion of over 50% overlap between peaks in two replicates using the bedtools *intersect* module (*-f 0.5*). The YTHDC1-associated R-loop sites were determined by the peaks that were exclusively present in two replicates of YTHDC1 KD samples, but not in the two replicates of control samples using the bedtools *intersect* module (*-f 0.5 -v*). Only the sites with at least 10 raw reads in all the replicates of at least 1 condition were included in the subsequent analyses. To calculate YTHDC1 CLIP read coverage over the defined m6A, m1A, or YTHDC1-associated R-loop sites, we first normalized each CLIP data to its total read depth and apply the contrast against its corresponding input RNA data (CLIP - Input). The read coverage was quantified by calculating the number of reads in each normalized CLIP dataset at each m6A, m1A, or YTHDC1-associated R-loop site using bedmap *--count* module. The final read coverage for each condition (control vs. MMS treatment) was determined by calculating the average read coverage from two independent replicates at each m6A, m1A, or YTHDC1-associated R-loop site.

### Quantification of R-loop enrichment

To determine total R-loop sites, we combined the identified R-loop sites from both the control and YTHDC1 knockdown samples as described above. We quantified R-loop enrichment by calculating the MapR read count at each R-loop site for the YTHDC1 KD and control samples, respectively. Only the sites with at least 10 raw reads in all the replicates of at least 1 condition were included in the subsequent analyses. To determine the R-loop sites with significant differences in enrichment between the YTHDC1 knockdown and control samples, we utilized differential read count analysis using the generalized linear models implemented in edgeR (glmQLFTest function) by first normalizing the MapR read count of each sample using the TMM method implemented in the edgeR package, followed by contrasting the YTHDC1 knockdown sample against the control sample after normalization to their respective input samples ((KD_GST-RNΔ-MNase_ - KD_GST-MNase_) - (Control_GST-RNΔ-MNase_ - Control_GST-MNase_)). We determined R-loop enrichment at a site to be increased or decreased based on whether the log fold change in MapR read counts in the YTHDC1 KD sample was greater than 1 or less than -1, respectively, in comparison to the control sample with an adjusted p-value of less than 0.05.

### icSHAPE analysis

The *in vivo* icShape deep sequencing data was obtained from Lu *et al*. (2016)^36^ (GEO: SRX1605093, SRX1605094, SRX1605089, and SRX1605090). The icShape reactivity was calculated as described in Lu *et al*. (2016)^36^ and Spitale *et al*. (2015)^48^. Briefly, the reads were aligned to the GENCODE Release 45 (GRCh38.p14) genome using Bowtie2 (version 2.4.1). The icSHAPE reverse transcriptase (RT) stops were isolated as the first nucleotide after the 5’end barcode. RT stops were used to calculated icSHAPE reactivity scores. The reactivity score was defined as the subtraction of background reverse transcription stops (DMSO libraries) from reverse transcription stops of the modified NAI-N3 libraries, and then adjusted by the background base density. The coverage profile of reactivity score at m1A sites +/- 2kb was plotted by deeptools computematrix (reference-point) and plotProfile modules using the reactivity scores obtained above.

### Statistical Analyses

All p-values related, except where indicated, were calculated by unpaired, two-tailed Student’s t-test. Cell viability assay p-values were determined by two-way ANOVA. IF intensity, comet assay, and R-loop quantitation p-values were determined by Mann-Whitney test. Sequencing analysis p-values were determined by nonparametric Wilcoxon matched-pairs signed rank test. All error bars represent the standard deviation of the mean, unless otherwise noted.

### Antibodies

All antibodies for this study are shown in Extended Data Table 2.

## Acknowledgments

We thank Hani Zaher, Eric Greer, Kenneth Murphy, and Andre Nussenzweig for suggestions during the course of these studies. We thank Pei-Chi Wei (Heidelberg University) and Martijn Luijsterberg (Leiden University Medical Center) for kindly sharing the S9.6 antibody and RPE-1 cell lines, respectively. We thank Ross Tomaino (Harvard Medical School Taplin Core) for proteomics analysis, as well as the Genome Technology Access Center (GTAC), and the Genome Engineering and iPSC (GEiC) Center at Washington University for sequencing service. This work was supported by the NIH (R01 CA193318 and P01 CA092584), an American Cancer Society Research Scholar award (RSG-18-156-01-DMC), the Barnard Foundation, Centene Corporation, and the Siteman Cancer Center.

## Author Contributions

N.T., J.O., R.R., J.R.B., and N.M. carried out cellular and biochemical experiments. H.S. and C-K.C. performed bioinformatic analysis and wrote the corresponding methods. M-S.T. performed protein purification of GFP-dRNH. E.A.P. performed protein purification of GST-MNase and GST-RNΔ-MNase. N.T. and N.M. wrote the manuscript with inputs from all other authors. N.M. supervised the project.

## Competing Interests

The authors declare no competing financial interests.

## Materials and correspondence

All requests for materials and correspondence should be made to N.T. and N.M.

## Extended Data Figure Legends

**Extended Data Fig. 1.**
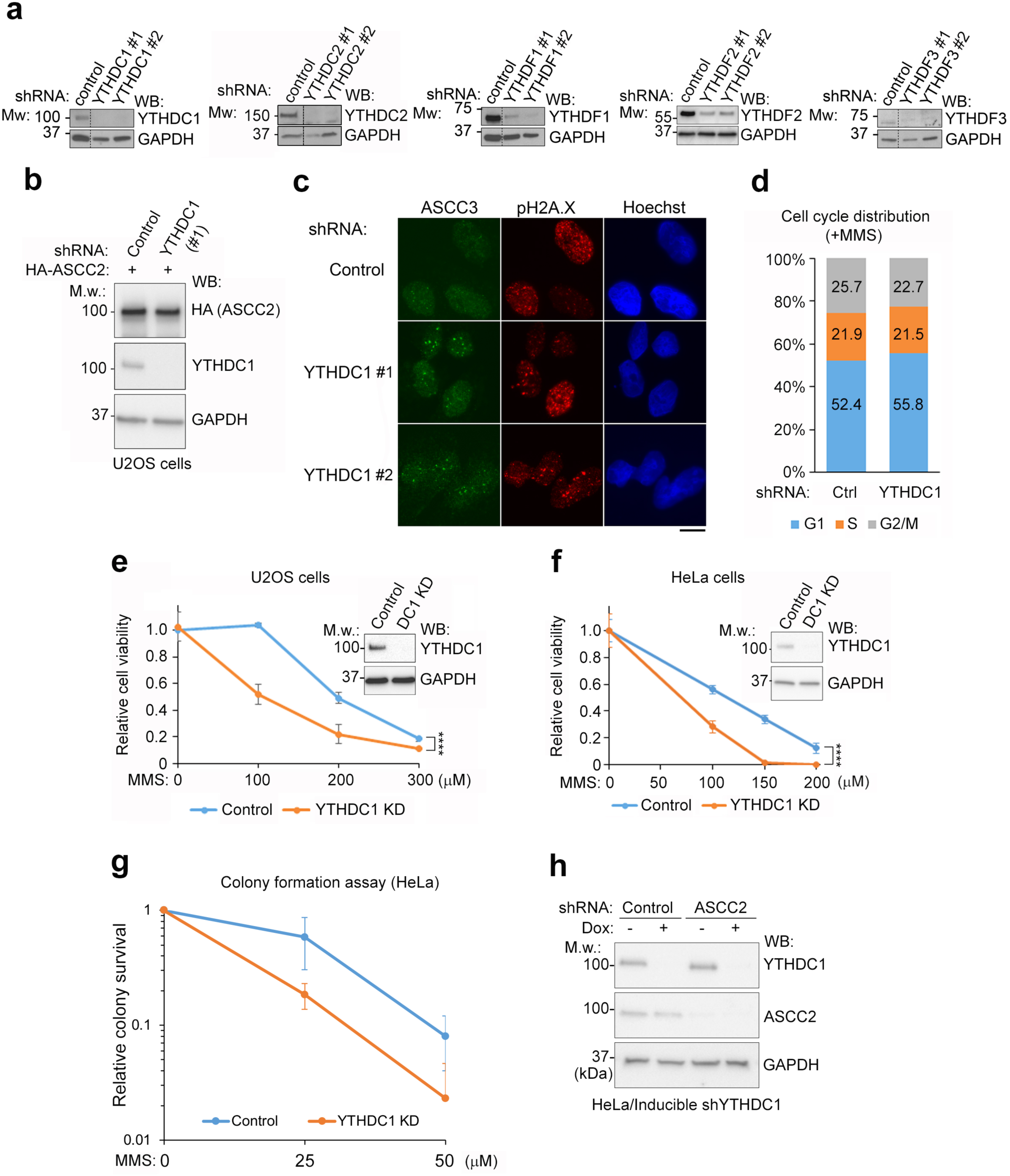
Characterization of YTHDC1 in the alkylation damage response. **a,** U2OS cells were infected with lentiviral shRNAs targeting individual human YTH proteins. Whole cell lysates were processed for western blotting with the indicated antibodies. **b**, Western blotting of lysates from HA-ASCC2 expressing U2OS cells infected with control or YTHDC1 lentiviral shRNA. **c,** Control and YTHDC1 shRNA expressing U2OS cells were treated with 0.5 mM MMS for six hours, then processed with immunofluorescence staining with indicated antibodies. Scale bar, 10 µm. **d**, Control and YTHDC1 shRNA expressing cells were treated with 0.5 mM MMS for 6 h, then processed with propidium iodide (PI) staining and flow cytometry for analyzing cell cycle distribution. **e, f,** MTS survival assays with U2OS (**e**) and HeLa (**f**) cells expressing inducible YTHDC1 shRNA with (YTHDC1 KD) or without (control) the induction of YTHDC1 shRNA using 0.5 µg/mL of doxycycline. Error bars indicate the mean ± SD (n = 5 technical repeats; ****p < 0.0001 by two-way ANOVA test). **g,** HeLa cells expressing inducible YTHDC1 shRNA with (YTHDC1 KD) or without (control) the induction of YTHDC1 shRNA were treated with the indicated doses of MMS for 24 hours. The cells were recovered with fresh media and incubated for another 14 days for colony formation. Relative survival level was indicated by normalizing the colony number of each sample to its corresponding control sample. Error bars indicate the mean ± SD (n = 3). **h**, HeLa cells expressing inducible YTHDC1 shRNA were infected with control and ASCC2 lentiviral shRNAs with or without the induction of YTHDC1 shRNA with 0.5 µg/mL of doxycycline. Cell lysates were then processed for western blotting with the indicated antibodies.

**Extended Data Fig. 2.**
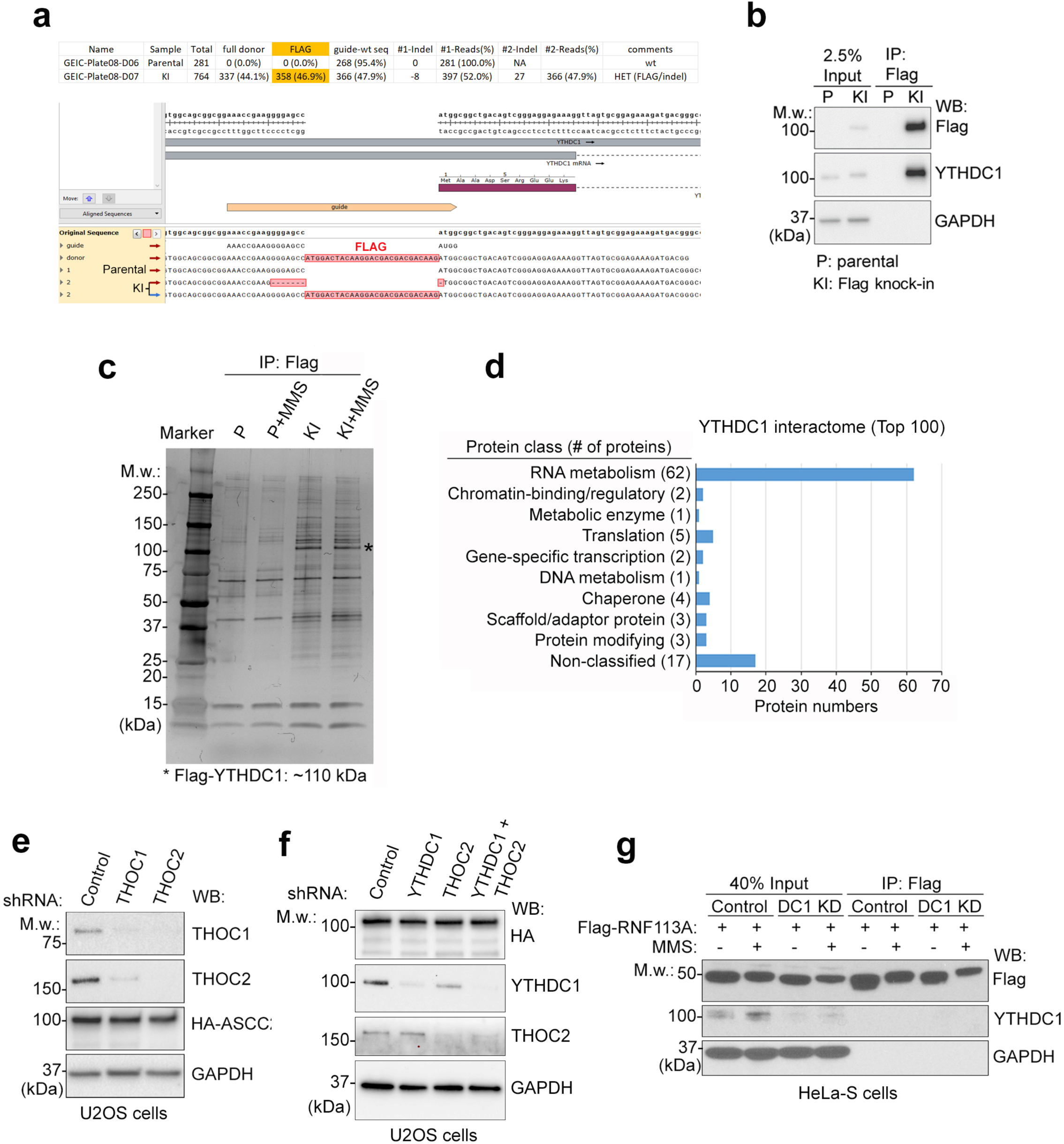
Identification of YTHDC1-interacting proteins and determining the function of THOC in alkylation responses. **a**, Verification of an endogenous Flag tag knock-in clone by deep sequencing. **b**, Parental (P) and endogenously Flag-tagged YTHDC1 (KI) HeLa-S cells were harvested for immunoprecipitation with anti-Flag (M2) resin. The input and immunoprecipitated materials were analyzed by western blot with the indicated antibodies. **c**, Silver staining of the YTHDC1 complex separated on a 4%-12% SDS-PAGE. The YTHDC1 complex was immunopurified from nuclear extracts prepared from parental (P) and endogenously Flag-tagged YTHDC1 (KI)-containing HeLa-S cells. **d**, Panther GO term analysis (protein classification) of the YTHDC1-interacting proteins using the top 100 most abundant interacting proteins. **e-f**, U2OS cells expressing HA-ASCC2 were infected with the indicated lentiviral shRNAs. Whole cell lysates were used for western blotting with the antibodies as shown. **g**, Flag-RNF113A expressing HeLa-S cells were infected with control and YTHDC1 shRNAs (indicated as DC1 KD). Cells with or without MMS treatment were harvested for RIP-MS analysis (see Fig. 1j). To confirm that the recovery of Flag-RNF113A was not affected by depleting YTHDC1, the protein level of immunocomplexes and the corresponding input samples were analyzed by western blotting with the indicated antibodies.

**Extended Data Fig. 3.**
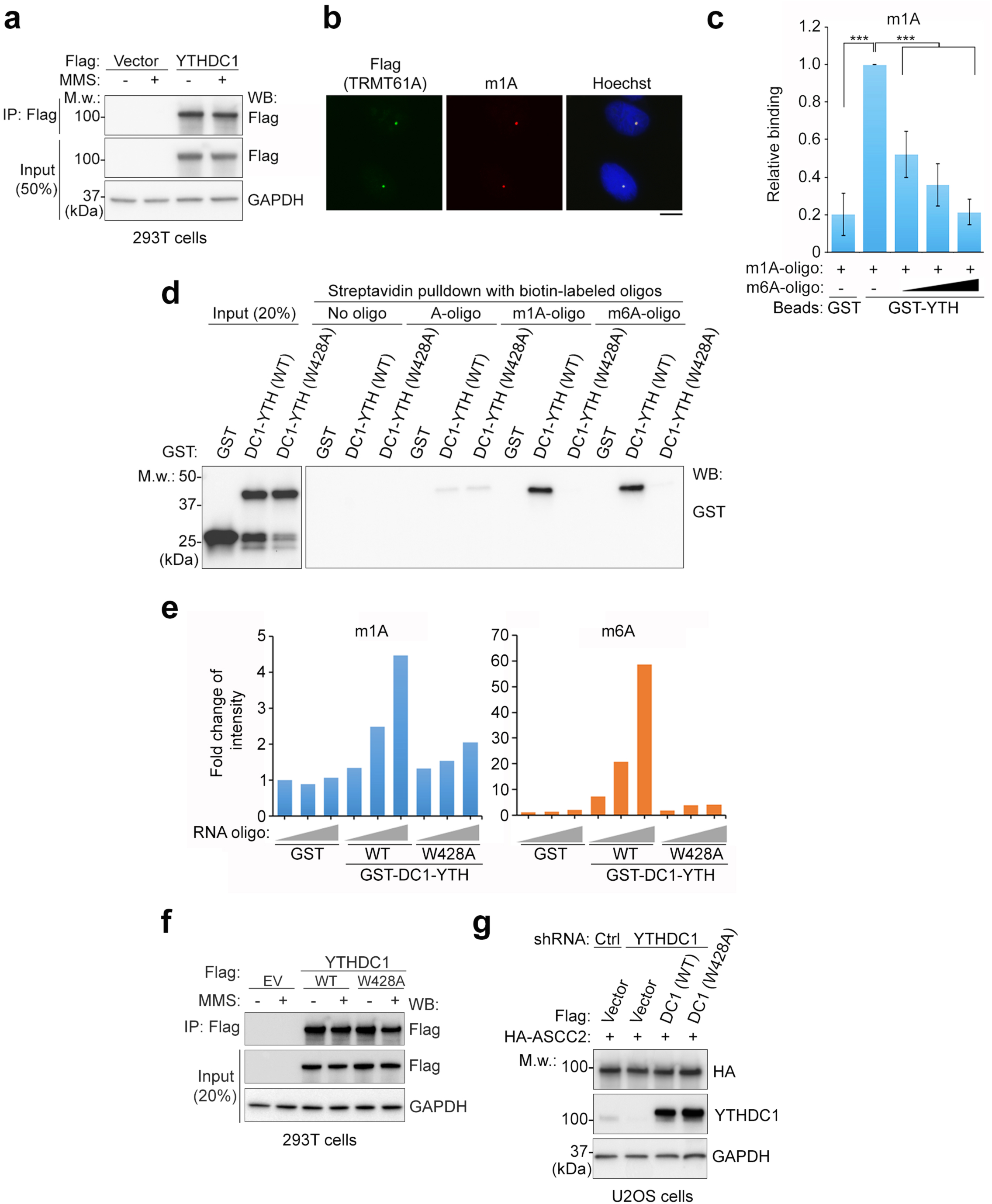
YTHDC1 interacts with m1A-containing RNAs via its YTH domain. **a,** 293T cells expressing empty Flag vector and Flag-YTHDC1 were harvested for RIP-MS (see **Fig. 2a**). The immunoprecipitates (IP) and the corresponding input samples were analyzed by western blotting with the indicated antibodies. **b,** The DD degron/Flag-tagged TRMT61A-LacI expressing U2OS 2-6-3 single locus reporter cells were incubated with 300 nM Shield1 and 1 µg/mL doxycycline for 24 hours, then processed for immunofluorescence staining with the indicated antibodies. **c,** GST recombinant proteins were incubated with m1A-containing oligonucleotides and increasing amount of m6A-containing oligonucleotidess, then pulled down using GST beads. The relative quantity of m1A recovered from each RNA pulldown was determined by quantitative LC-MS/MS (n = 3 independent experiments; error bars indicate ± SD of the mean; ***p < 0.001 by Student’s t test). **d,** Biotin-labeled RNA oligonucleotides (unmodified A, m1A, or m6A) were incubated with wildtype or W428A GST-YTH domain of YTHDC1; bound material was then pulled down using streptavidin beads. The bound proteins were analyzed by western blotting using anti-GST antibody. **e,** Purified GST, wildtype and W428A GST-YTH domain of YTHDC1 (0.2 µM of each) were incubated with increasing amounts of m1A- or m6A-containing oligonucleotides (0.1, 0.2, or 0.5 µM); the bound material as then pulled down using GST beads. The quantity of m1A- or m6A-containing oligonucleotide bound was determined by LC-MS/MS. **f,** 293T cells expressing empty Flag vector and the indicated Flag-YTHDC1 were harvested for RIP-MS (see Fig. 2c). The immunoprecipitates (IP) and the corresponding input material were analyzed by western blotting with the indicated antibodies. **g,** HA-ASCC2 expressing U2OS cells were infected with the indicated lentiviral shRNAs, as well as empty Flag vector or the indicated Flag-YTHDC1 expression vectors. Whole cell lysates were used for western blotting using the indicated antibodies.

**Extended Data Fig. 4.**
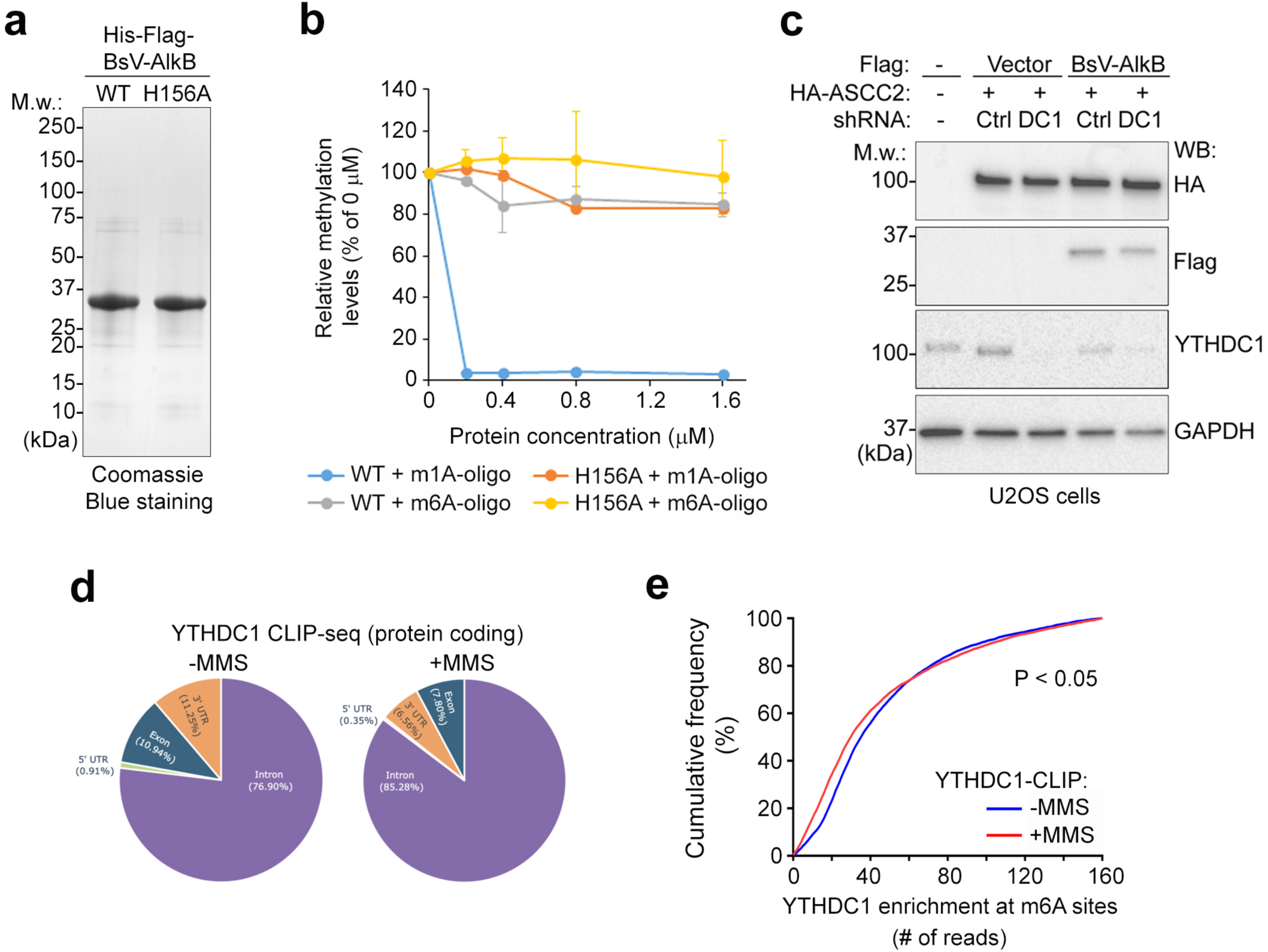
*In vitro* demethylase assay with BsV-AlkB and YTHDC1 CLIP-seq analysis. **a**, Coomassie blue staining of the recombinant wildtype (WT) and catalytically inactive (H156A) His-tagged BsV-AlkB purified from *E. coli*. **b**, *In vitro* demethylase assays on m1A- and m6A-containing RNA oligonucleotides (0.4 µM of each) with increasing concentration of WT and H156A BsV-AlkB proteins as shown. Reactions were quantified using LC-MS/MS and shown as a percentage of no protein control (0 µM). Error bars indicate ± SD of the mean (n = 3 technical replicates). **c,** HA-ASCC2 expressing U2OS cells were infected with the indicated lentiviral shRNAs along with empty Flag or the indicated Flag-BsV-AlkB expression vectors. Whole cell lysates were used for western blot analysis using the indicated antibodies. **d-e**, CLIP-seq analysis of YTHDC1 with (+) or without (-) MMS-induced damage. High confidence hits within specific regions mapped to protein coding RNAs are shown in (**d**). Cumulative frequency in percentage (%) of YTHDC1 enrichment at m6A sites from YTHDC1 CLIP-seq with or without MMS treatment was shown in (**e**) (bin size = 1; maximum bin capped at 160). *p < 0.05 by using a nonparametric Wilcoxon matched-pairs signed rank test.

**Extended Data Fig. 5.**
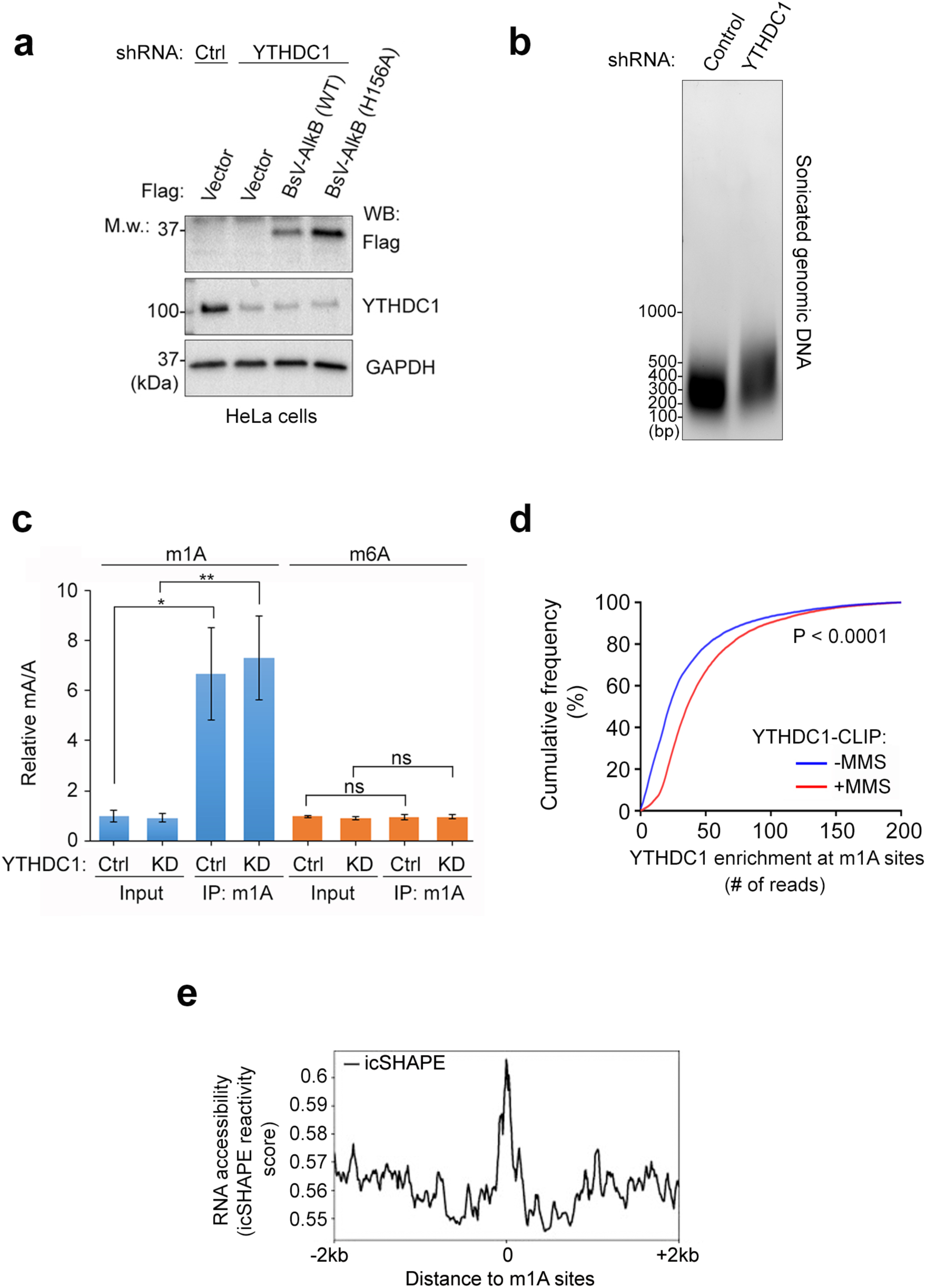
BsV-AlkB expression and analysis of m1A DNA immunoprecipitation. **a**, HeLa cells were infected with the indicated lentiviral shRNAs and co-expressed with empty Flag vector and the indicated Flag-BsV-AlkB expression vectors. Protein expression was analyzed by western blotting with the indicated antibodies. **b**-**c**, HeLa cells expressing control and YTHDC1 shRNA were treating with 2 mM MMS for one hour, then processed for nuclear isolation and genomic DNA extraction (see Fig. 4e). DNA samples were fragmented by sonication and analyzed as shown (**b**). This was followed by immunoprecipitation with m1A antibody and LC-MS/MS analysis (**c**). Levels of m1A and m6A were quantitated by LC-MS/MS (normalized to the level of A). The result is shown as the mean ± SD (n = 3 independent experiments; *p < 0.05, **p < 0.01 by Student’s t test; ns, non-significant). **d**, Cumulative frequency in percentage (%) if YTHDC1 enrichment at m1A sites from YTHDC1 CLIP-seq with or without MMS treatment (bin size = 1; maximum bin capped at 200). ****p < 0.0001 of MMS-treated samples compared to non-MMS-treated samples using a nonparametric Wilcoxon matched-pairs signed rank test. **e**, icSHAPE analysis revealed that higher RNA accessibility (icShape reactivity) was observed at the center of m1A sites.

**Extended Data Fig. 6.**
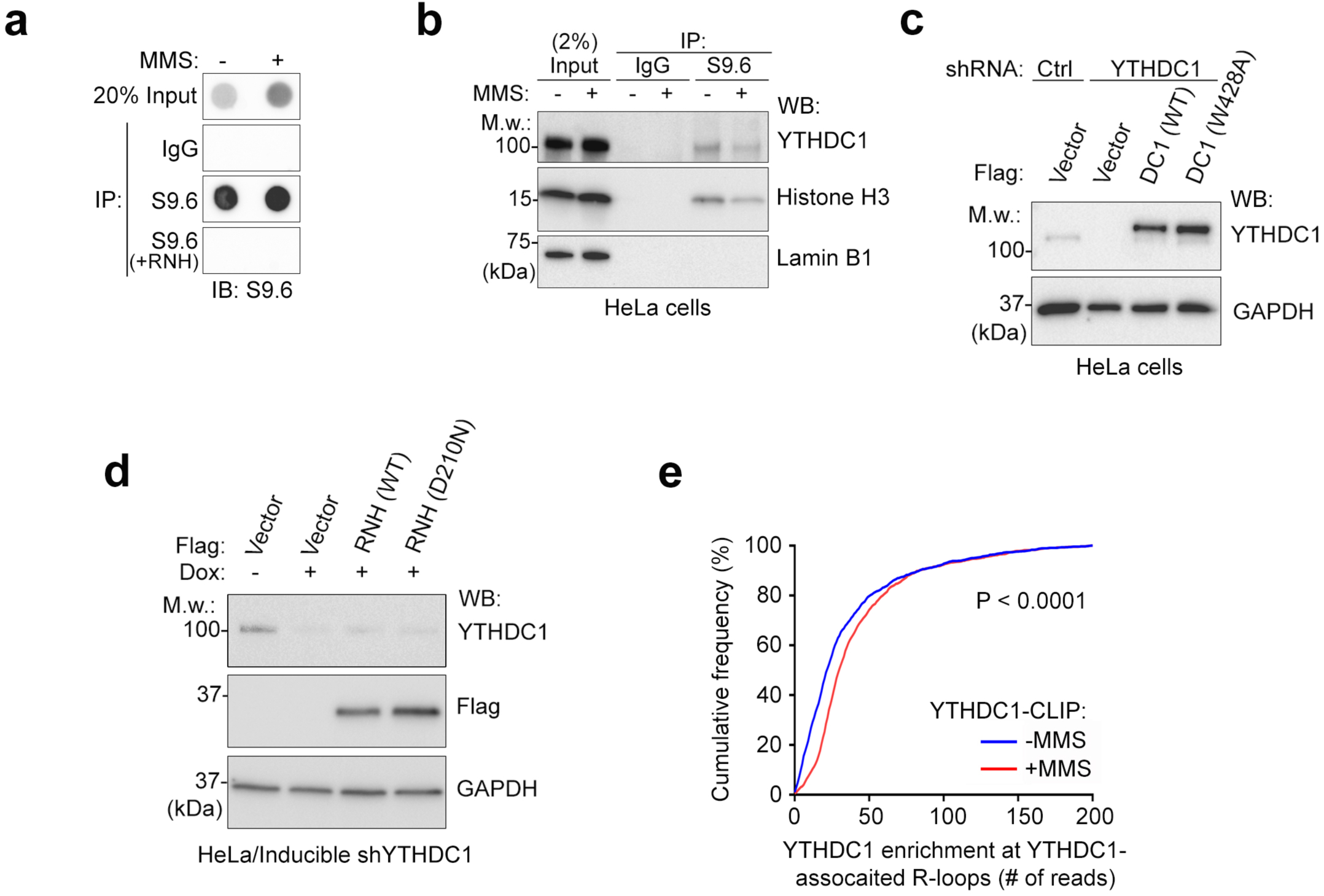
YTHDC1 is associated with R-loops and genome-wide R-loop analysis. **a**, Genomic DNAs were purified from HeLa cells treating with (+) or without (-) 2 mM MMS for one hour. DNA samples pre-treated with or without RNaseH1 were immunoprecipitated using S9.6 (RNA:DNA hybrid) antibody or normal mouse IgG as control. Recovered material from the immunocomplexes were subjected to dot blot with S9.6 antibody. **b**, Nuclear extracts were prepared from HeLa cells treated as in (**a**), then immunoprecipitated with S9.6 antibody or mouse IgG. The IP and input material was analyzed by western blotting and probed using the indicated antibodies. **c**, Control or YTHDC1 shRNA-expressing HeLa cells were co-transduced with vectors expressing empty Flag or the indicated Flag-YTHDC1 expression vectors. Protein expression was analyzed by western blotting with the indicated antibodies. **d**, HeLa cells expressing doxycycline-inducible YTHDC1 shRNA were co-transduced with the indicated RNaseH1 (RNH) expression vectors with or without the induction of YTHDC1 shRNA by 0.5 µg/mL of doxycycline. Cell lysates were then processed for western blotting with indicated antibodies. **e**, Cumulative frequency in percentage (%) of YTHDC1 enrichment at YTHDC1-associated R-loops from YTHDC1 CLIP-seq with or without MMS treatment (bin size = 1; maximum bin capped at 200). ****p < 0.0001 of MMS-treated samples compared to non-MMS-treated samples using a nonparametric Wilcoxon matched-pairs signed rank test.

**Extended Data Fig. 7.**
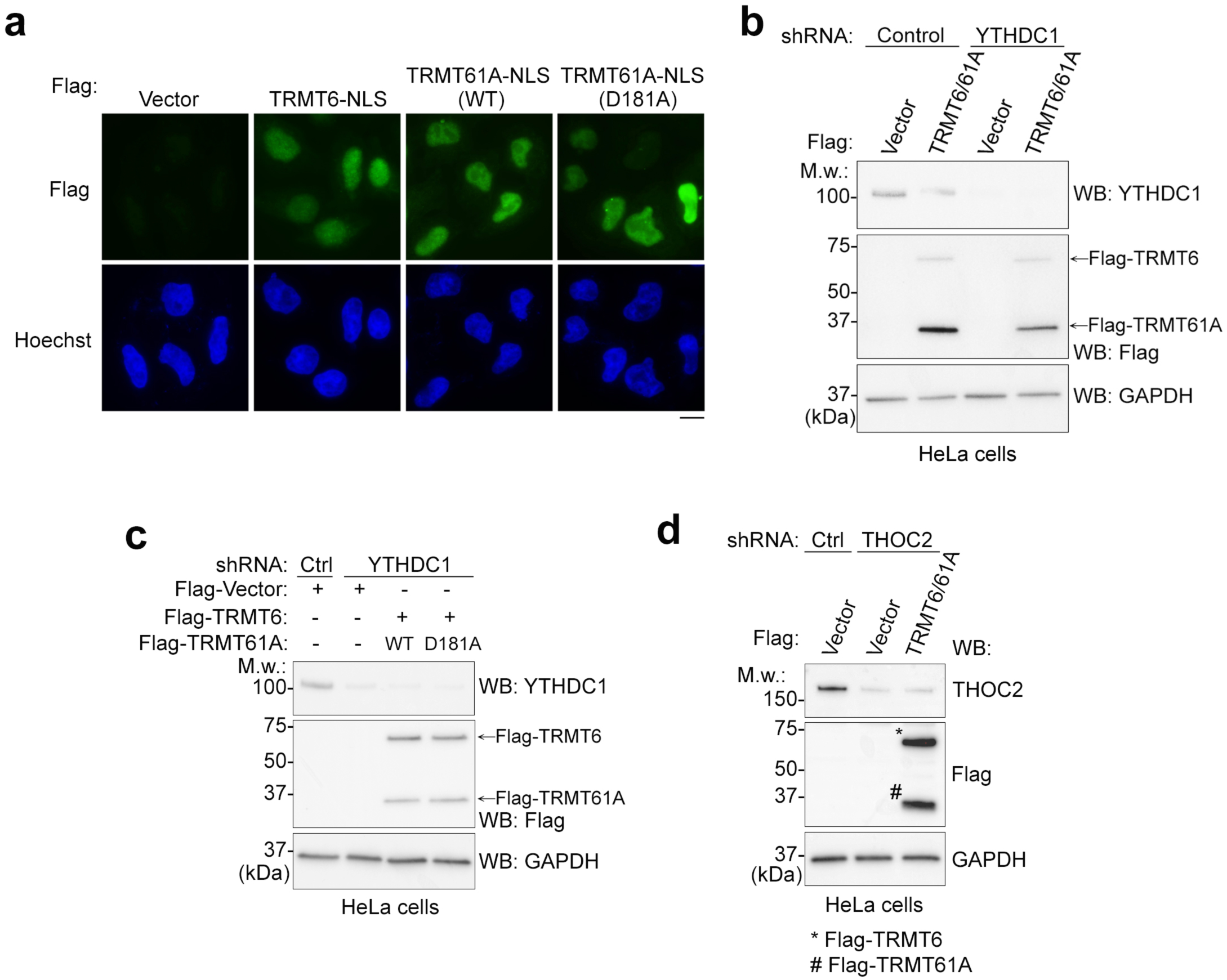
Expression of the NLS-tagged TRMT61A and TRMT6 in cells. **a,** HeLa cells were infected with empty Flag vector or the indicated TRMT-NLS lentiviral expression vectors. After 48 hours, cells were fixed and immunofluorescence staining with Flag antibody was performed. Scale bar, 10 µm. **b-c,** Control and YTHDC1 shRNA-expressing HeLa cells were co-transduced with empty Flag vector or the indicated Flag-TRMT6-NLS or Flag-TRMT61A-NLS expression vectors. Protein expression was analyzed by western blotting with the indicated antibodies. **d**, Control and THOC2 shRNA-expressing HeLa cells were co-transduced with empty Flag vector or the NLS tagged Flag-TRMT expression vectors. Protein expression was analyzed by western blotting with the indicated antibodies.

## Notes

### Competing Interest Statement

The authors have declared no competing interest.

